# Validated Antimalarial Drug Target Discovery using Genome-Scale Metabolic Modelling

**DOI:** 10.1101/2025.03.26.645484

**Authors:** Supannee Taweechai, Francis Isidore Garcia Totañes, David Westhead, Clara Herrera-Arozamena, Richard Foster, Glenn A. McConkey

## Abstract

Given the rapid resistance of *Plasmodium falciparum* to antimalarial drugs, there is a continual need for new treatments. A genome-scale metabolic (GSM) model was developed with integrated omic and constraint-based, experimental flux-balance data to predict genes essential for *P. falciparum* growth as drug targets. We selected the highly ranked *P. falciparum* UMP-CMP kinase (UCK) to test its necessity and the ability to inhibit growth with inhibitors. Conditional deletion mutants using the DiCre recombinase system, generated by CRISPR-Cas genome editing, exhibited defective asexual growth and stage-specific developmental arrest. Based on *in silico* and *in vitro* screening, inhibitors were identified that are selective for *P. falciparum* UCK and exhibit antiparasitic activity. This study, for the first time, shows assertions from a GSM model identifying novel, validated, “druggable” targets. These findings show a role for GSM models in antimalarial drug discovery and identify *P. falciparum* UCK as a novel, valid malaria drug target.

## 1. Introduction

Malaria is a significant global health burden, focussed in tropical and subtropical regions, with 249 million malaria infections globally and circa 608,000 malaria-related deaths(1). Chemotherapy, the main approach to treatment, is restricted. For example, artemisinin-based combination therapies (ACTs), currently recommended and widely used for malaria treatment in endemic countries worldwide, exhibited decreased susceptibility in the Greater Mekong subregion in 2009 with delayed parasite clearance (2–5). Hence, the rapid and regular arising of resistance necessitates continuous antimalarial drug discovery to new targets for control of infection.

With the constant need for new antimalarial drugs, methods have been developed for identifying drug targets on the genomic level. Identifying genes whose loss of function results in a severe fitness defect as ‘essential’ in genome-wide knockouts is a first step in drug target validation. Publication of *Plasmodium falciparum* saturation mutagenesis, that was recently extended to *Plasmodium knowlesi* with a considerably higher number of mutations, classified a large collection of genes into essential and non-essential based on subtraction of mutations that do not affect growth (6, 7). This binary classification of genes may generate false positive essential genes (e.g. due to factors such as growth requirements under different environmental conditions or non-random transposon insertions due to chromatin structure) and multiple transposons inserted in a genome may also yield compensatory effects. Hence, directed mutagenesis of single genes is needed for validation. Genome-scale metabolic (GSM) models (aka GEMs) integrate omics and experimental data to identify targets that are most susceptible to growth inhibition based on flux-balance analysis. Constraint-based models can prioritise enzymes that are most likely to be “druggable” (8). A stage-specific Plasmodium GSM model has fostered prioritisation of drug targets based on blood stages of infection (9). Their utility in asserting and prioritising drug targets has been reviewed extensively but have not followed progression of drug target validation from modelling to target validation and druggability. In this study, a constraint-based model of *P. falciparum* was developed with prioritised assertion of drug targets and a top-ranked gene was characterised and validated by development of a directed, inducible knockout of the gene. The target was assessed as drug target in biochemical assays with inhibitors and tested for parasite growth inhibition.

The research herein investigates the assertion that UMP-CMP kinase (UCK) is a viable drug target from development of a *P. falciparum* GSM model. The study validates the necessity for *P. falciparum* UCK (PfUCK), investigates its potential as an antimalarial drug target by employing CRISPR-Cas generated inducible mutants, and assesses its druggability for future drug development. This includes biochemical analysis and development of a spectrophotometric assay for PfUCK that facilitates future high-throughput inhibitor screening.

## 2. Materials and methods

### 2.1. Metabolic model construction

The three malaria metabolic models that were merged together in this study were those developed by Forth (10), Plata (11) and Huthmacher (12). As the three models utilized different ontological formats for metabolite (i.e., species) and reaction IDs, these IDs were initially converted to SEED format using database files of reaction and species IDs in various ontological formats obtained from metanetx.org (13). the Forth model (iTF143) was used as the base model (also referred to as the minimal model). The Huthmacher and the Plata models served as the source models, i.e., models from which reactions were collected and added into the minimal model. Reactions from the source models must satisfy the following criteria to be added into the minimal model:

1. The reaction has enzyme commission classification and gene association data.
2. At least one species in the reaction is in the minimal model.

To ensure accuracy of comparison between reactions, the reaction equations were compared instead of the reaction IDs. Reactions from the source models that satisfy the two criteria were removed from the list of source model reactions and were added into the minimal model. As new reactions were being added to the minimal model, new species from these reactions were also added. Thus, it was possible that reactions that initially satisfied the first criteria but not the second could now have a common species in the growing minimal model. The remaining reactions from the source models were repeatedly assessed until no new reaction could be added into the model.

### 2.2. In vitro flux measurements

Continuous *P. falciparum* 3D7 in vitro cultures were grown in RPMI 1640 growth medium (Life Technologies, UK) supplemented with 5% (w/v) Albumax II (Gibco, USA), 0.01% (w/v) hypoxanthine (Sigma, USA) and 0.1% (v/v) gentamicin at 5% haematocrit (O+ blood obtained from the National Blood Service of the NHS Blood and Transplant in Seacroft, Leeds) in a 37°C incubator at 1% oxygen, 3% carbon dioxide and 96% nitrogen gas mixture. Cultures were synchronised with 5% sorbitol (14). Synchronised cultures with a total volume of 18 ml at 1% parasitaemia and 5% haematocrit were placed in non-vented 75 cm^2^ tissue culture flasks (Nunclon). Red blood cells at 5% haematocrit were used as control. One millilitre of culture was collected at time 0, and every 6 hours until 48 hours post synchronisation. The collected sample was placed in a 1.5 ml microcentrifuge tube and centrifuged at 3000 rpm for 2 minutes. The spent media was collected and stored at -80°C prior to analysis. Three biological replicates were analyzed.

A glucose assay on the spent media was performed using a glucose oxidase-peroxidase format assay kit (Megazyme). Amino acid concentrations in the spent media were determined using an Ultimate 3000 High-Performance Liquid Chromatography (HPLC) system (Dionex, UK). A ramp gradient reverse-phase chromatography was done with an Acclaim 120 C18 (Dionex, |UK) 100 mm x 2.1 mm column (3 μm particle size, 120 Å pore size) stationary phase. Eluent A (10 mM Na_2_HPO_4_, 10 mM Na_2_HB_4_O_7_ · 10 H_2_O, 0.5 mM NaN_3_, pH 8.2) and Eluent B (45% methanol (v/v) 45% acetonitrile (v/v) in water) constituted the mobile phase (Supplementary Table 3).

Samples were derivatised manually within 10 minutes prior to chromatography. The derivatisation procedure was done in a 1.5 ml microcentrifuge tube at room temperature by adding the following reagents in this particular order:

1. 300 μl of borate buffer (0.1 M Na_2_HB_4_O_7_ · 10H_2_O, pH 10.2)
2. 15 μl of o-phthaldialdehyde (OPA) (75 mM OPA, 225 mM 3-mercaptopropionic acid in 0.1 M borate buffer, pH 10.2)
3. 3 μl of sample (mixed 5 times using a Gilson pipette set at 300 μl)
4. 6 μl 9-fluorenylmethoxycarbonyl chloride (FMOC) solution (2.5 mg/mL FMOC in acetonitrile) (mixed 5 times using a Gilson pipette set at 300 μl)
5. 42 μl phosphoric acid solution (15 μl/ml 85% phosphoric acid in Eluent A)

Detection of derivatised amino acids was through UV absorbance at 338 nm at 10.0 Hz data collection rate (from 0 to 55 minutes).

Spent media for both infected and uninfected (control) samples were assayed for glucose and amino acid concentrations as detailed above. The net metabolite flux from time 0 (*t*_0_) to time n (*t*_𝑛_) was calculated using the formula:

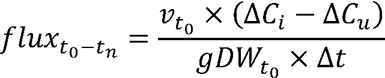

where:

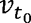 = volume at *t*_0_, in litres
Δ𝐶_𝑖_ = change in metabolite concentration in infected culture from *t*_0_ − *t*_𝑛_, in mM
Δ𝐶_𝑢_ = change in metabolite concentration in uninfected culture from *t*_0_ − *t*_𝑛_, in mM
𝑔𝐷𝑊_*t*0_ = gram dry weight of parasite at *t*_0_
Δ*t* = change in time (*t*_𝑛_ − *t*_0_), in hours

The parasite mass 𝑔𝐷𝑊_*t*0_ was determined using the calculated number of parasites based on the parasitaemia, haematocrit and the volume at *t*_0_:

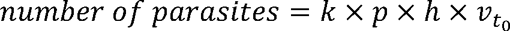

where:

𝑘 = haematocrit constant 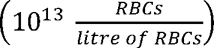
𝑝 = parasitaemia at t_0_
ℎ = haematocrit at t_0_
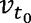 = volume at t_0_, in litres

In this case, it was assumed for simplicity that each infected RBC contained only one parasite. The total parasite mass (𝑔𝐷𝑊) was then calculated by multiplying the number of parasites by the mass per parasite experimentally measured previously 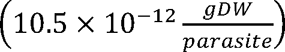 (15). Flux calculations were done to represent the three main blood stage forms: the change in concentration from *t*_6_to *t*_18_ was used to calculate the early to mid-ring stage flux, from *t*_30_ to *t*_36_ for the late trophozoite and from *t*_42_ to *t*_48_ for the late schizont stage. Stage-specific average flux values ± standard deviations were used as upper and lower boundary flux constraints, respectively, for the corresponding boundary transport reaction. A positive flux represents entry of metabolites into the model while a negative flux represents movement of metabolites into the external environment.

### 2.3. Model simulations

To mimic *in vitro* consumption of nucleoside/bases from the media, the boundary flux of adenine, adenosine, guanine and xanthine were set to zero as these are not present in Albumax II supplemented RPMI media. This allowed the model to consume hypoxanthine for purine metabolism from the external environment.

Single gene knockout simulation was also done using COBRApy. A given gene knockout that produced less than 95% of the optimal biomass was identified as growth limiting whereas a knockout that produced a zero biomass flux was considered lethal. Predicted essential genes were compared against published data on experimentally validated essential genes/reactions in *Plasmodium* (i.e., gold standard list). Furthermore, gene essentiality data from the *Plasmodium* Genetic Modification (PlasmoGEM) database were also used to obtain essential genes that were added to the gold standard list. PlasmoGEM, developed as part of the initiative of the Malaria Programme at the Wellcome Trust Sanger Institute, is a database that holds prepublication phenotypic data on more than 2,000 *Plasmodium berghei* genes (16, 17). Updated *P. berghei* gene IDs along with phenotypic data were downloaded from the PlasmoGEM database and orthologues of these genes in *P. falciparum* were identified through PlasmoDB (18). For *P. falciparum* genes that are orthologous to more than one *P. berghei* gene, only those with consistent gene essentiality information were noted (e.g., all orthologous *P. berghei* genes must be essential). The hypergeometric p-value was calculated to identify any significant enrichment of experimentally validated essential genes/reactions in the model predictions.

### 2.4. Plasmid construction

A repair plasmid, designated pL6-UCK_loxP-sgRNA4-native E3-I3l, was created based on the plasmid pL6 eGFP, generously provided by Prof. Jose-Juan Lopez-Rubio from the Biology of Host-Parasite Interactions Unit at Institut Pasteur, Paris, France. This repair plasmid contains both a single guide RNA (sgRNA) and a donor DNA template essential for homology-directed repair, where the h*DHFR* gene is flanked by two homology regions (HR1 and HR2). To construct the repair plasmid, the GFP regions (GFP 5’ and GFP 3’) of the pL6 eGFP plasmid were replaced with the homology regions (HR1 and HR2) of the gene of interest (GOI). Subsequently, the targeting oligonucleotides of sgRNA were incorporated at the *Btg*ZI adapter region following the method outlined by (19).

In the initial step, sgRNA4 (CATAATGAATTAAATGACCA), identified using the online Benchling tool, was introduced into the pL6-eGFP plasmid utilizing the NEBuilder HiFi DNA assembly kit (New England BioLabs Inc, USA). Following this, the donor DNA sequence, encompassing the homology-directed repair 1 (HDR1) domain—nucleotides 113–1108 of the genomic DNA sequence—was inserted. In this arrangement, the upstream *loxP* sequence was positioned within intron 2 (between nucleotide 684 and 720), followed by the recoded PfUCK gene (nucleotides 1109-1765), the downstream *loxP* site, the hDHFR gene serving as the selectable marker, and the 3’ UTR sequence forming the homology-directed repair 2 (HDR2). This process was facilitated using the NEBuilder HiFi DNA assembly kit (NEB Inc, USA) to integrate these elements into the plasmid pL6 eGFP.

The repair plasmid pL6-UCK_loxP-sgRNA4-native E3-ΔI3 was derived from the plasmid pL6-UCK-sgRNA4-native E3-I3 by eliminating the native intron 3 sequence. To accomplish this, two sets of primers were employed. The HDR1 forward primer (5’-CTTTCCGCGGGGAGGACTAGTACATAAATAGAACTAATGGTT-3’) and the native Exon3_reverse primer (5’-CGTTGTTTATACAATCCTCA-3’) were used to amplify the fragment containing nucleotides 113–864 from the plasmid pL6-UCK-sgRNA4-native E3-I3. Subsequently, the exon4 forward primer (5’-TGAGGATTGTATAAACAACGGTAAAATCGTGCCGG-3’) and the native exon4 reverse primer (5’-TTTTTTTTCAACACCAGATTCACCCTGGTCGTT-3’) were used to amplify the fragment containing nucleotides 1026-1108 from the genomic DNA, with the exon4 forward primer designed to exclude the intron 3 sequence. Consequently, these amplified fragments were inserted into the plasmid pL6-UCK-sgRNA4-native E3-I3, with no alterations made to the recoded PfUCK gene (nucleotides 1109-1765), the downstream *loxP* site, the hDHFR gene, or the 3’ UTR sequence, which serve as the homology-directed repair 2 (HDR2).

### 2.5. P. falciparum transfection and cloning

The DiCre-expressing parasite strain B11 (20) was cultured in supplemented RPMI1640 growth medium as above. The parasite cultures were incubated horizontally at 37°C with a low oxygen gas mixture as previously described (21). For synchronization, ring stage parasites were generated by treating with 5% D-sorbitol for 10 minutes. To enrich the late stages (trophozoite and schizont) of *P. falciparum* parasites, a method based on the sedimentation behaviour of late-stage parasites in gelatin solution, described by (22) and (23) with minor modifications, was employed. Approximately 7%–8% parasitaemia of asexual stages (mostly trophozoite stage parasites) were pelleted using centrifugation at 250 x g for 10 minutes, and the supernatant was discarded. The pellet was then resuspended in prewarmed media at a 3:1 ratio (medium: pellet), followed by the addition of two volumes of gelatin solution (ZeptoGel; Gentaur). After gentle mixing, the solution was incubated at 37°C for 30 minutes. The supernatant containing the trophozoites and schizonts was transferred into a new culture flask, ready for transfection of DNA-loaded erythrocytes.

The transfection protocol, followed the preloading of erythrocytes as described by (24). Positive selection drugs, namely 1 nM WR99210 (gifted by Jacobus Pharmaceuticals) and 250 nM DSM265 (synthesized by the Chemistry Department at the University of Leeds), were applied after 48 hours and continued for four days. Upon detection of parasites via thin blood smearing, negative selection drug 40 µM 5-fluorocytosine (5-FC) was applied for five days to eliminate parasites carrying episomal plasmids. Subsequently, parasites were cloned using a limiting dilution method, aiming for a calculated dilution of 0.5 parasites per well with 3% haematocrit. Each well of 96-well flat-bottom plates contained 200 µL of the diluted parasite culture and was incubated in a humidified sealed chamber with low oxygen at 37°C. Parasite maintenance included changing the medium every other day, with daily flushing of cultures with low oxygen gas. Fresh RBCs were replenished every five days for parasite reinvasion. Samples were collected between days 10–14 for thin blood smear analysis. Subsequently, selected positive parasite clones were transferred to T25-flasks and cultured in complete RPMI 1640 media with 5% haematocrit.

### 2.6. Conditional knockout induction

DiCre-driven *loxP site* recombination for gene function analysis was induced by treating tightly synchronised ring-stage of transgenic parasite clones with 50 nM RAP (Stratech Scientific Ltd, Ely) or mock treatment using 0.1% v/v DMSO for 24 hours. Transgenic parasite control F5, carrying a single *loxP* site, was used to control for unexcised parasites. At 42 hours post-RAP treatment, 200 µL of parasite cultures were collected for gDNA extraction by the microwave irradiation method (25). Next, the truncated Pf*UCK* was assessed by PCR analysis of parasite gDNA using primers U6 (TCTACCTTTTAAAAAATAGCAAGCA) and P1 (CAAGTATATATTTTGTTTCTATAAATTGATATCTTA).

### 2.7. Parasite growth assay

Parasite growth was determined in samples collected on days 0, 2, and 4 post-RAP treatment with parasitaemia measured via fluorescence-activated cell sorting (FACS) as previously described (26). Parasite samples were stained with SYBR Green I nucleic acid gel stain (Thermo Fisher Scientific, UK) by the following procedure: Initially, 100 µL of parasite cultures were centrifuged at 800 x g for 1 minute. Subsequently, the cell pellets were washed with 500 µL of phosphate-buffered saline (PBS) and centrifuged again. The parasite samples were then fixed with 100 µL of 4% paraformaldehyde (Thermo Fisher Scientific, UK) and 0.1% glutaraldehyde (Sigma-Aldrich, USA) in PBS for 1 hour at 4°C, followed by another PBS wash. Next, the cells were stained with 1:5000 SYBR Green I in PBS for 1 hour at 37°C, washed with PBS, and resuspended in 100 µL of PBS. Finally, 10 µL of stained cells were diluted into 1 mL of PBS in an Eppendorf tube and analysed using a CytoFLEX S Flow Cytometer with a 525/40 BP filter to determine the proportion of SYBR Green-stained cells per 100,000 cells. Statistical analysis of the growth assay was conducted using GraphPad Prism 9 with a two-way ANOVA analysis (Bonferroni’s multiple comparison test), with results indicating statistical significance for P<0.05.

### 2.8. Gene expression quantitation

Total RNA was isolated from parasite pellets utilising the Direct-zol RNA MiniPrep kit (Zymo Research, USA) in accordance with the manufacturer’s instructions. Subsequently, genomic DNA (gDNA) was eliminated from RNA samples using the Turbo DNA-free kit (Invitrogen). Then, 120 ng of DNA-free RNA samples served as the template for cDNA synthesis employing the Maxima H Minus First Strand cDNA Synthesis kit (Thermo Fisher Scientific), with oligo (dT)18 acting as the reverse transcription primer. For RT-qPCR reactions, the Brilliant III Ultra-Fast QPCR Master Mix (Agilent Technologies, USA) was utilised. Amplification of the PfUCK gene was carried out using the C3F2 forward primer (5’-TGAGGATTGTATAAACAACGGTAAAATCGTGCCGG-3’) and C3F2 reverse primer (5’-TTTTTTTTCAACACCAGATTCACCCTGGTCGTT-3’), while the small subunit rRNA (SSU rRNA) reference gene was amplified using the ssu_q_forward (5’-CGAACGAGATCTTAACCTGC-3’) and ssu_q_reverse (5’-CACACTGTTCCTCTAAGAAGC-3’) primers. RT-qPCR was initiated with an initial denaturation step at 95°C for 2 minutes, followed by 40 cycles of denaturation at 95°C for 30 seconds, annealing at 65°C for 30 seconds, and extension at 68°C for 30 seconds, carried out in a C100 Thermo Cycler (Bio-Rad, USA) with default settings. Thermal melt assay revealed single amplicons of the anticipated size. Relative changes in gene expression were determined using the 2^-ΔΔCT^ method (27). Statistical analysis using two-tailed Student’s *t-*tests was performed to assess the significance of changes in mRNA levels as unpaired samples.

### 2.9. UCK protein expression and purification

The native PfUCK protein has a hydrophobic region spanning amino acid residues 8 to 23 as predicted by the Malaria Secretory Signal Predictions (MalSig) program. The hydrophobic core at the protein’s N-terminus may cause poor expression and low solubility in *E. coli*. Therefore, a truncated gene was created where the ORF began at amino acid residue 24–371, encoding a 348 amino acid protein. Subsequently, the optimised codons for recombinant expression of the truncated Pf*UCK* was generated using the program OPTIMIZER (28), then was commercially synthesized and cloned into the plasmid pUC57 (GenScript). Using a pair of primers (5’-TGCCGCGCGGCAGCCATATGGAAAACTTCTACCTGCTG-3’ and 5’-TCGAGTGCGGCCGCAAGCTTTTACATGTTGGAAAACG-3’), the truncated gene was amplified from the plasmid pUC57 and subsequently sub-cloned into the pET28a expression vector via NdeI and HindIII cloning sites. The pET28a plasmid containing PfUCK was transformed into *E. coli* BL21 (DE3) Rosetta cells, which were then selected and cultured for subsequent protein purification. Briefly, protein expression was induced with isopropyl-D-thiogalactoside, incubated at 16°C with shaking at 200 rpm for 20 hours, and the cell pellet resuspended in binding buffer (50 mM Tris [pH 7.6], 300 mM NaCl, 20 mM imidazole, 5% glycerol, and 0.075% β-mercaptoethanol), with cell disruption using an Avestin C3 Cell Disrupter. Cell debris was removed and the UCK purified by Ni-NTA affinity purification. The protein was eluted from the column using a linear gradient of imidazole (20–500 nM) in elution buffer (50 mM Tris [pH 7.6], 300 mM NaCl, 5% glycerol, and 0.075% β-mercaptoethanol). PfUCK-containing fractions were combined and concentrated before assessing purity via 12% SDS-PAGE. The concentrated protein was then stored at -70°C in a storage buffer (25 mM Tris-HCl, 50 mM KCl, and 20% glycerol; pH 7.0).

Recombinant human UMP-CMP kinase (h*UCK*) was constructed by amplifying its ORF from an available commercial plasmid (CMPK1 [NM_016308] Human Tagged ORF Clone–RC204856; OriGene) using primers hUCK forward (5’-TGCCGCGCGGCAGCCATATGATGCTGAGCCGC-3’) and reverse (5’-TCGAGTGCGGCCGCAAGCTTTTAGCCTTCCTTGTCAAAAAT-3’). The gene was cloned into the pET28a expression plasmid and hUCK expressed in induced BL21 (DE3) RIL cells similar to PfUCK. The protein was purified with a HisPur Ni-NTA Spin column, and elution with 250 mM imidazole in elution buffer (50 mM Tris-HCl, 750 mM NaCl, and 10% glycerol; pH 7.5). The protein was further purified using a His GraviTrap TALON column. The buffers employed were: binding buffer (50 mM potassium phosphate [KP], 100 mM NaCl, and 10 % glycerol; pH 8.0), washing buffer (50 mM KP, 500 mM NaCl, 10 mM imidazole, and 10 % glycerol; pH 8.0), and elution buffer (50 mM Tris-HCl, 750 mM NaCl, 250 mM imidazole, and 10% glycerol; pH 7.5). The purified protein was desalted using a PD-10 column (Sephadex G25M; Cytiva) resulting in a storage buffer (25 mM Tris-HCl, 50 mM KCl, and 20% glycerol; pH 7.0). Purity of the recombinant protein was verified via SDS-PAGE (Mini-Protein TGX Precast Gels; Bio-Rad), and its concentration was determined using a Bradford protein assay (Bio-Rad).

### 2.10. UCK characterization

Enzyme activity was assessed using an indirect spectrophotometric assay that measured NADH absorbance at 340 nm coupled to pyruvate kinase/lactate dehydrogenase. The reaction mixture contained 50 mM Tris-HCl (pH 7.6), 50 mM KCl, 10 mM MgCl_2_, 5 mM ATP, 0.2 mM NADH, 1 mM phosphoenolpyruvate, 10 mM DTT, pyruvate kinase (10 U/mL), lactate dehydrogenase (15 U/mL), and purified UCK (OD340/min of 0.04–0.06). The reaction was initiated by adding 1 mM of the substrate (CMP, UMP, or dCMP) and the assay was performed at 25°C for 10 minutes in the absence or the presence of UCK. The decrease in absorbance was monitored using a UV/Visible spectrophotometer (Ultrospec 2100 Pro).

UCK’s kinetic properties were determined by varying nucleoside monophosphate (CMP, UMP, and dCMP) concentrations. Then, a reaction rate was calculated from the decrease in absorbance at 340 nm (ε_NADH_=6.22×10^-3^ M^-1^ cm^-1^) compared to the control reaction in which UCK was absent. The maximum reaction rate (*V*_max_) and Michaelis constant (*K*_m_) values were calculated using the Michaelis–Menten equation (*v*=*V*_max_[*S*] / *K*_m_+[*S*]), where *v* is the reaction rate and [*S*] is the substrate concentration.

### 2.11. Inhibition constant (K_i_) determination

The 1 mL reaction assay consisted of 50 mM Tris-HCl (pH 7.6), 50 mM KCl, 10 mM MgCl2, 5 mM ATP, 0.2 mM NADH, 1 mM phosphoenolpyruvate, 10 mM DTT, pyruvate kinase (10 U/mL), lactate dehydrogenase (15 U/mL), and appropriate amounts of purified UCK (yielding an OD340/min of 0.04–0.06) along with varying concentrations of the inhibitor. The final concentration of DMSO in the assay reaction was maintained at 1%. Compounds were initially screened for inhibition of recombinant PfUCK enzyme activity using a 10-fold serial dilution in dimethyl sulfoxide (DMSO). Active compounds were then further tested in a two-fold serial dilution to determine their *K*_i_ value. The enzyme assay was as described above, with initiation by adding 1 mM CMP. As inhibitors were obtained from a virtual screening using the model structure of PfUCK as the template, *K*_i_ values for each inhibitor were determined using nonlinear regression curve fitting in Prism software based on a competitive equation utilising CMP’s Km value for calculation. Using the competitive equation here is based on an assumption that those inhibitors bind the active site of a protein. Compounds active against PfUCK were also tested for their *K*_i_ values against hUCK using the same protocol as for PfUCK.

### 2.12. Antiplasmodial activity (IC_50_) determination

*P. falciparum 3D7* parasites underwent two synchronization cycles with D-sorbitol before assay day. Parasitaemia and haematocrit were adjusted to 0.5% and 3%, respectively. Synchronized ring parasites were cultured in 96-well plates in complete RPMI medium with varying inhibitor concentrations. Inhibitor stock solutions, mostly at 100 mM, were prepared in DMSO. However, some compounds had 50 mM stock solutions due to solubility limitations. Triplicate wells were prepared for each compound. Negative controls included wells with uninfected red cells (3% haematocrit) in complete RPMI medium.

Culture plates were incubated in a chamber with controlled gas conditions at 37°C. After 48 hours, parasites were quantified using a fluorescence-based assay with SYBR Green dye as previously described (21). Lysis buffer containing SYBR Green was added to each well, and plates were incubated at room temperature for 45 minutes and protected from the light. Fluorescence was measured using a microplate reader (POLARstar OPTIMA, BMG Labtech) with excitation at 485 nm and detection at 520 nm. Background fluorescence of uninfected red cells was subtracted, and dose-response curves were plotted against inhibitor concentration. IC_50_ values were determined using nonlinear regression curve fitting in GraphPad Prism software.

## 3. Results

### 3.1 Development of a combined genome-scale metabolic model

Three malaria metabolic models developed by Forth (10), Plata (11) and Huthmacher (12) were combined to develop a highly curated genome-scale metabolic model supported by experimentally derived metabolite data. Having the fewest dead-end metabolites and reactions, the Forth model was used as the base model and reactions from the two other models were added in an iterative manner (i.e., reassessing after every addition of reaction). This method ensured no additional dead ends were incorporated into the final model while maximizing the total number of reactions contributed by the Plata and Huthmacher models.

The final model, iFT342, has a total of 342 genes, 551 reactions and 560 metabolites (Supplementary data 1). The model includes five compartments: the apicoplast, cytosol, mitochondria, vacuole and the external compartment. Out of the 551 reactions in the model, 371 (67.3%) are gene-associated, most of which are intracellular reactions. Most of the non-gene associated reactions are transport reactions (16.5%) and boundary reactions (13.4%), while only 2.5% are intracellular reactions. The model has a biomass reaction that represents the consumption of different components necessary for parasite growth. The stoichiometry of the biomass equation was derived using experimentally quantified biomass macromolecular components (DNA, RNA and proteins) together with published data on the proportions of different subcomponents of these macromolecules (29). There are 106 boundary metabolites and 454 intracellular metabolites. All 560 metabolites in the model have chemical formulas, and 530 have additional metabolite attributes (i.e., PubChem ID (30), IUPAC International Chemical Identifier (InChI) keys (31) and canonical Simplified Molecular-Input Line-Entry System (SMILES) (32). Figure 1 shows the distribution of reactions based on the gene association, compartment, enzyme commission (EC) classification and subsystem involvement.

**Figure 1.**
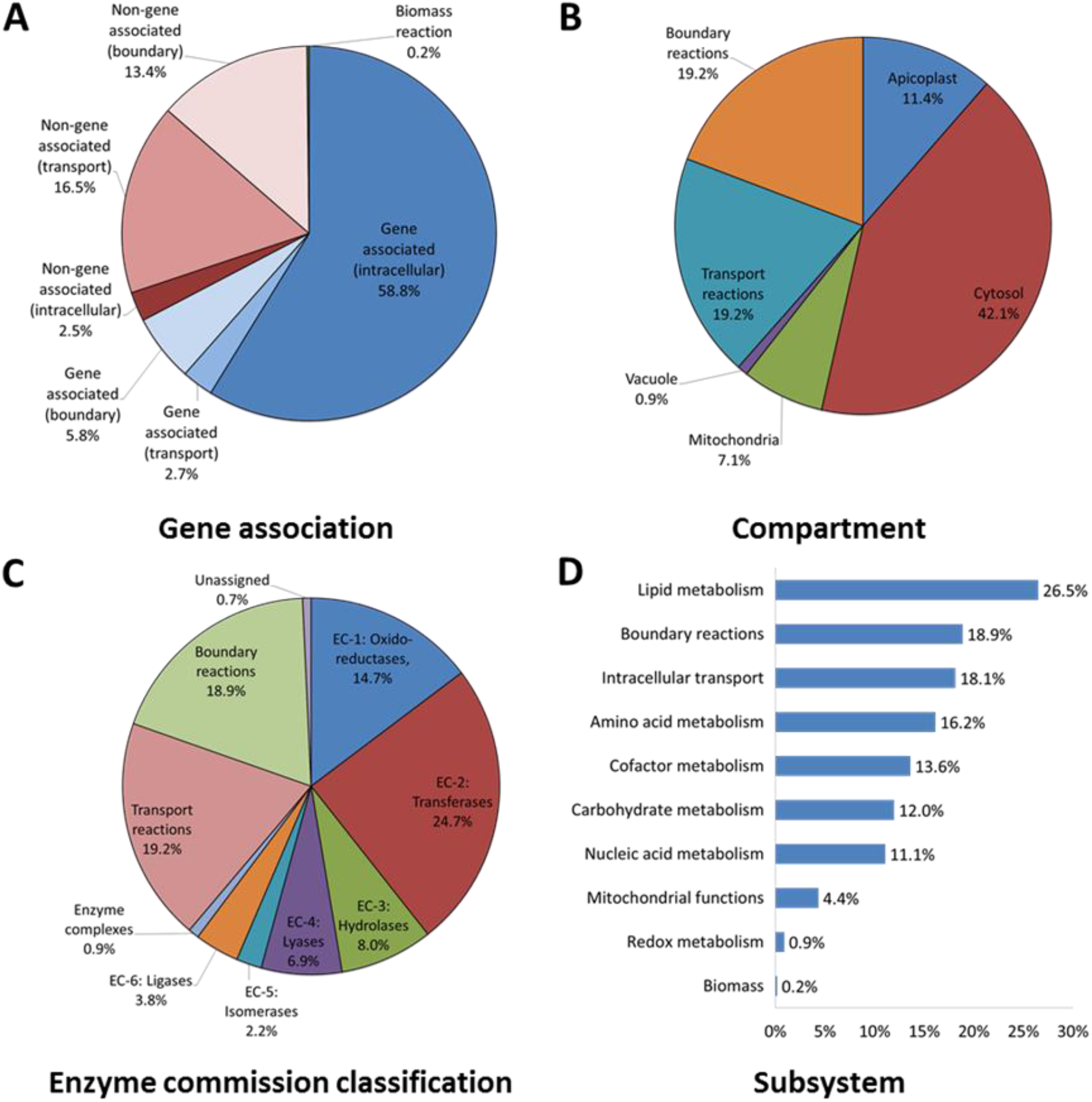
Reactions in the iFT342 model grouped by (A) gene association, (B) compartment, (C) enzyme commission classification and (D) subsystem involvement.

In vitro flux of glucose and 18 amino acids were measured in highly synchronised in vitro cultured *P. falciparum* 3D7 at three different time points to represent the early to mid-ring trophozoite stages, late trophozoite stages and late schizont stages (Supplementary Table 1). In addition, measurements of lactate in spent media from previous experimental data conducted in our facility were incorporated into the model (10). These flux values were incorporated into the model to generate stage-specific models. The standard deviation above and below the calculated mean flux were assigned as the upper and lower bound constraints of the corresponding boundary reaction in the model, respectively.

### 3.2. Genome-scale modelling asserts a possible antimalarial target

Using COBRApy (33), single gene knockout simulations were done to identify essential genes in the schizont stage model (Supplementary Figure 1). Out of 342 genes in the model, 87 genes were predicted to be essential: 47 gene knockouts were predicted to be lethal (i.e., biomass solution = 0), while 40 were growth-limiting (i.e., biomass solution < 95% optimal biomass solution). This was compared to a list of essential genes which is a combination of *P. falciparum* genes defined as essential based on chemical or genetic validation in published literature (i.e., gold standard) and a set genes orthologous to essential *P. berghei* genes that were defined based on being unable to generate gene knockouts for those genes (PlasmoGEM) (16) resulting in a list composed of a total of 1,257 genes (Supplementary Table 2). Overall, 58 (66.7%) of the predicted essential genes are in the list of experimentally validated essential genes, giving an enrichment of 1.51 compared to total percentage of experimentally validated essential genes in the whole model (151 out of 342, 44.2%; hypergeometric p-value = 6.41 x 10^-7^). Interestingly, *all* the predicted growth-limiting essential genes are in the experimentally validated essential gene list (enrichment of 2.26; hypergeometric p-value = 2.29 x 10^-16^). The model also predicted 18 novel *P. falciparum* gene-associated targets that are not in the gold standard list (Table 1). These assert potential antimalarial drug targets. As the majority of antimalarial drugs inhibit enzymes conserved in humans but targeting non-conserved residues, these have potential as drug targets.

**Table 1.**
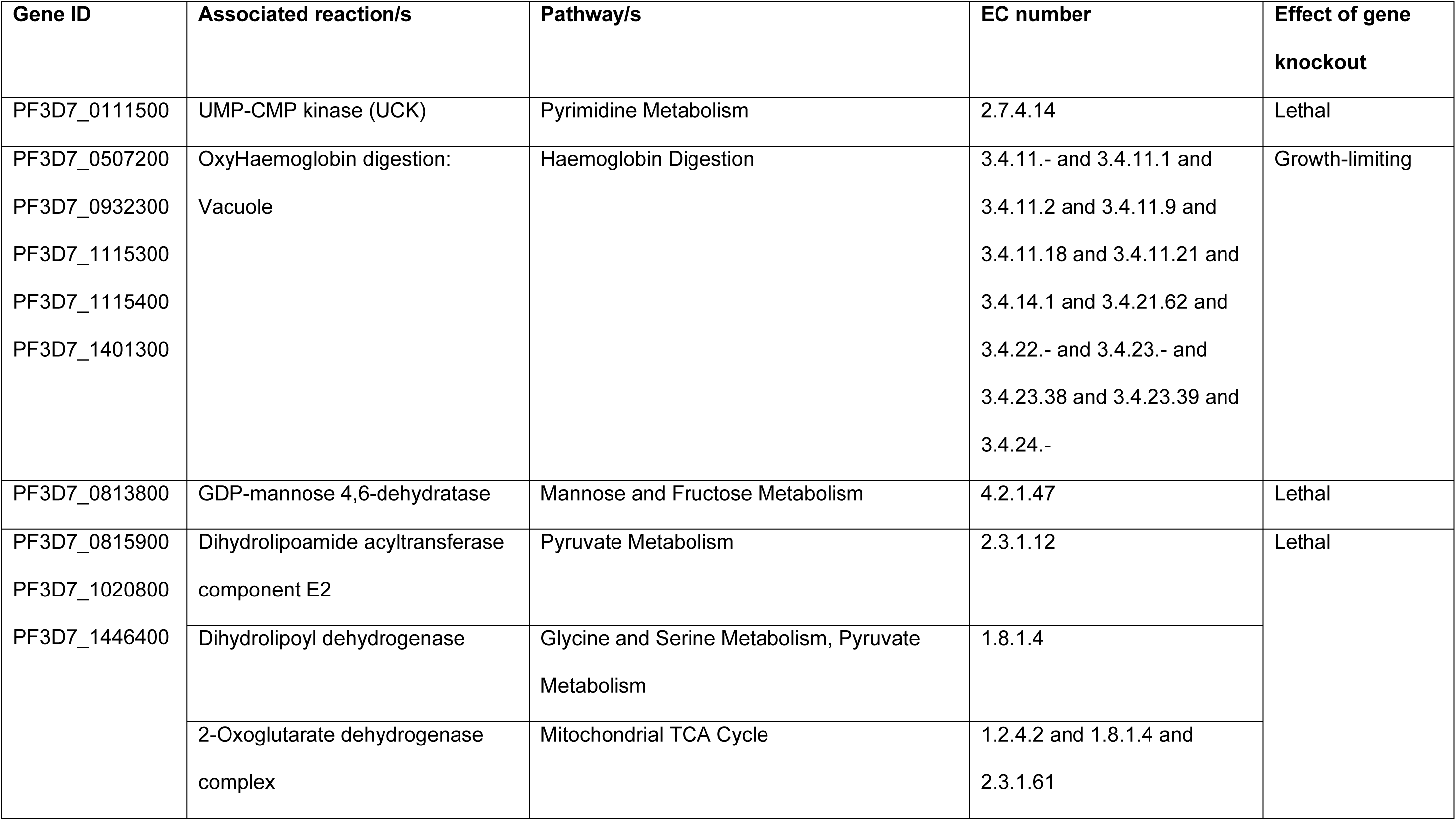
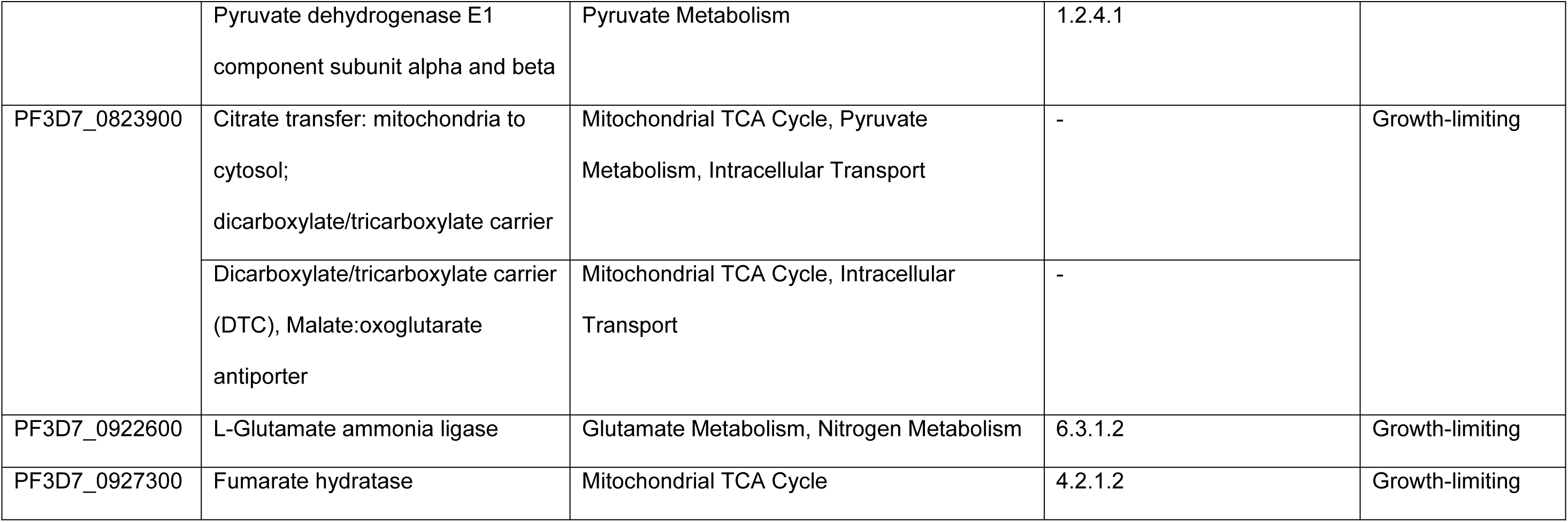
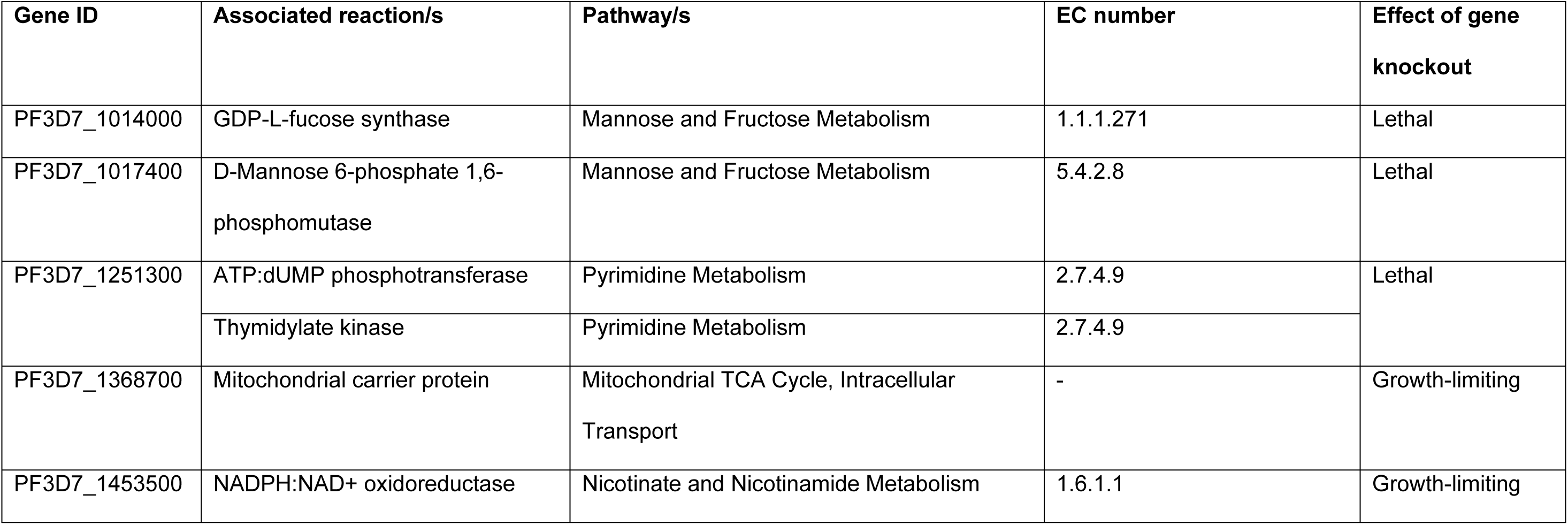
List of novel gene targets and their associated reactions as predicted by single gene knockout using the schizont stage model.

| Gene ID | Associated reaction/s | Pathway/s | EC number | Effect of gene knockout |
| --- | --- | --- | --- | --- |
| PF3D7_0111500 | UMP-CMP kinase (UCK) | Pyrimidine Metabolism | 2.7.4.14 | Lethal |
| PF3D7_0507200<br>PF3D7_0932300<br>PF3D7_1115300<br>PF3D7_1115400<br>PF3D7_1401300 | OxyHaemoglobin digestion:<br>Vacuole | Haemoglobin Digestion | 3.4.11.- and 3.4.11.1 and<br>3.4.11.2 and 3.4.11.9 and<br>3.4.11.18 and 3.4.11.21 and<br>3.4.14.1 and 3.4.21.62 and<br>3.4.22.- and 3.4.23.- and<br>3.4.23.38 and 3.4.23.39 and<br>3.4.24.- | Growth-limiting |
| PF3D7_0813800 | GDP-mannose 4,6-dehydratase | Mannose and Fructose Metabolism | 4.2.1.47 | Lethal |
| PF3D7_0815900<br>PF3D7_1020800<br>PF3D7_1446400 | Dihydrolipoamide acyltransferase<br>component E2<br>Dihydrolipoyl dehydrogenase<br>2-Oxoglutarate dehydrogenase<br>complex | Pyruvate Metabolism<br>Glycine and Serine Metabolism, Pyruvate<br>Metabolism<br>Mitochondrial TCA Cycle | 2.3.1.12<br>1.8.1.4<br>1.2.4.2 and 1.8.1.4 and<br>2.3.1.61 | Lethal |
|  | Pyruvate dehydrogenase E1 component subunit alpha and beta | Pyruvate Metabolism | 1.2.4.1 |  |
| PF3D7_0823900 | Citrate transfer: mitochondria to cytosol;<br>dicarboxylate/tricarboxylate carrier | Mitochondrial TCA Cycle, Pyruvate Metabolism, Intracellular Transport | - | Growth-limiting |
|  | Dicarboxylate/tricarboxylate carrier (DTC), Malate:oxoglutarate antiporter | Mitochondrial TCA Cycle, Intracellular Transport | - |  |
| PF3D7_0922600 | L-Glutamate ammonia ligase | Glutamate Metabolism, Nitrogen Metabolism | 6.3.1.2 | Growth-limiting |
| PF3D7_0927300 | Fumarate hydratase | Mitochondrial TCA Cycle | 4.2.1.2 | Growth-limiting |
| PF3D7_1014000 | GDP-L-fucose synthase | Mannose and Fructose Metabolism | 1.1.1.271 | Lethal |
| PF3D7_1017400 | D-Mannose 6-phosphate 1,6-phosphomutase | Mannose and Fructose Metabolism | 5.4.2.8 | Lethal |
| PF3D7_1251300 | ATP:dUMP phosphotransferase | Pyrimidine Metabolism | 2.7.4.9 | Lethal |
|  | Thymidylate kinase | Pyrimidine Metabolism | 2.7.4.9 |  |
| PF3D7_1368700 | Mitochondrial carrier protein | Mitochondrial TCA Cycle, Intracellular Transport | - | Growth-limiting |
| PF3D7_1453500 | NADPH:NAD<sup>+</sup> oxidoreductase | Nicotinate and Nicotinamide Metabolism | 1.6.1.1 | Growth-limiting |

The protein sequences of the predicted novel targets were aligned against proteins with known interacting compounds (e.g., inhibitors, cofactors, substrate) in the DrugBank database (34). A total of ten novel targets aligned with protein sequences in the DrugBank database, with e values ranging from 3.75 x 10^-133^ to 3.46 x 10^-7^. Out of the ten, only two sequences were associated with known inhibitors. Table 2 shows the two novel targets that aligned against protein sequences in the DrugBank database with known inhibitors. Gemcitabine, which interrupts DNA synthesis through phosphorylation by UMP-CMP kinase, has been shown to be efficiently transported by the parasite’s equilibrative nucleoside transporter (PfENT1) (35). In tests for in vitro activity against *P. falciparum* 3D7, the drug inhibited parasite growth in a stage-specific manner with an IC_50_ of 2.14 ± 0.76 μM (Supplementary Figure 2) (36).

**Table 2.** Novel targets with known FDA-approved inhibitors.

| Gene ID | Gene name | E value | Drug name | Drug group | Drug class |
| --- | --- | --- | --- | --- | --- |
| PF3D7_0111500 | UMP-CMP kinase, putative | 1.08E-24 | Gemcitabine | approved | Anti-metabolite, anti-cancer |
| PF3D7_0815900 | Dihydrolipoyl dehydrogenase, apicoplast (aLipDH) | 1.24E-25 | Carmustine | approved | Anti-cancer |

Modelling predicts that UCK plays a crucial role in *P. falciparum* pyrimidine metabolism; converting uridine monophosphate (UMP), cytidine monophosphate (CMP), and deoxycytidine monophosphate (dCMP) into their respective diphosphate forms (UDP, CDP, and dCDP). These diphosphate forms are further metabolized into triphosphates (UTP, CTP, and dCTP), essential for DNA and RNA synthesis (37). Notably, *P. falciparum* relies solely on the *de novo* pathway for pyrimidine nucleotide biosynthesis, unlike human cells that utilize both salvage and *de novo* pathways (38). Additionally, UCK’s medicinal importance lies in its involvement in activating prodrugs like gemcitabine, troxacitabine, and lamivudine, used for treating cancer and viral infections (39–45).

### 3.3. Strategy for generation of inducible UCK knockout mutants

As attempts to recover viable parasites with disruptions in the Pf*UCK* coding region in *P. falciparum* were unsuccessful (unpublished observation), we developed a strategy for generation of inducible *UCK* knockout mutants. Two repair plasmids with *loxP* sites inserted in introns, namely pL6-UCK_loxP-sgRNA4-native E3-I3 and pL6-UCK_loxP-sgRNA4-native E3-ΔI3 were designed that permit inducible deletion of the UCK enzyme’s active site upon induction (Figure 2). Both plasmids were designed to induce a modified parasite, with an upstream *loxP* site strategically positioned in intron 2 and a downstream *loxP* site located at the end of the *UCK* gene. The plasmid pL6-UCK_loxP-sgRNA4-native E3-I3 retained exon 3 and intron 3 in their native sequence, while pL6-UCK_loxP-sgRNA4-native E3-ΔI3 was derived by eliminating the native intron 3 sequence from pL6-UCK-sgRNA4-native E3-I3.

**Figure 2.**
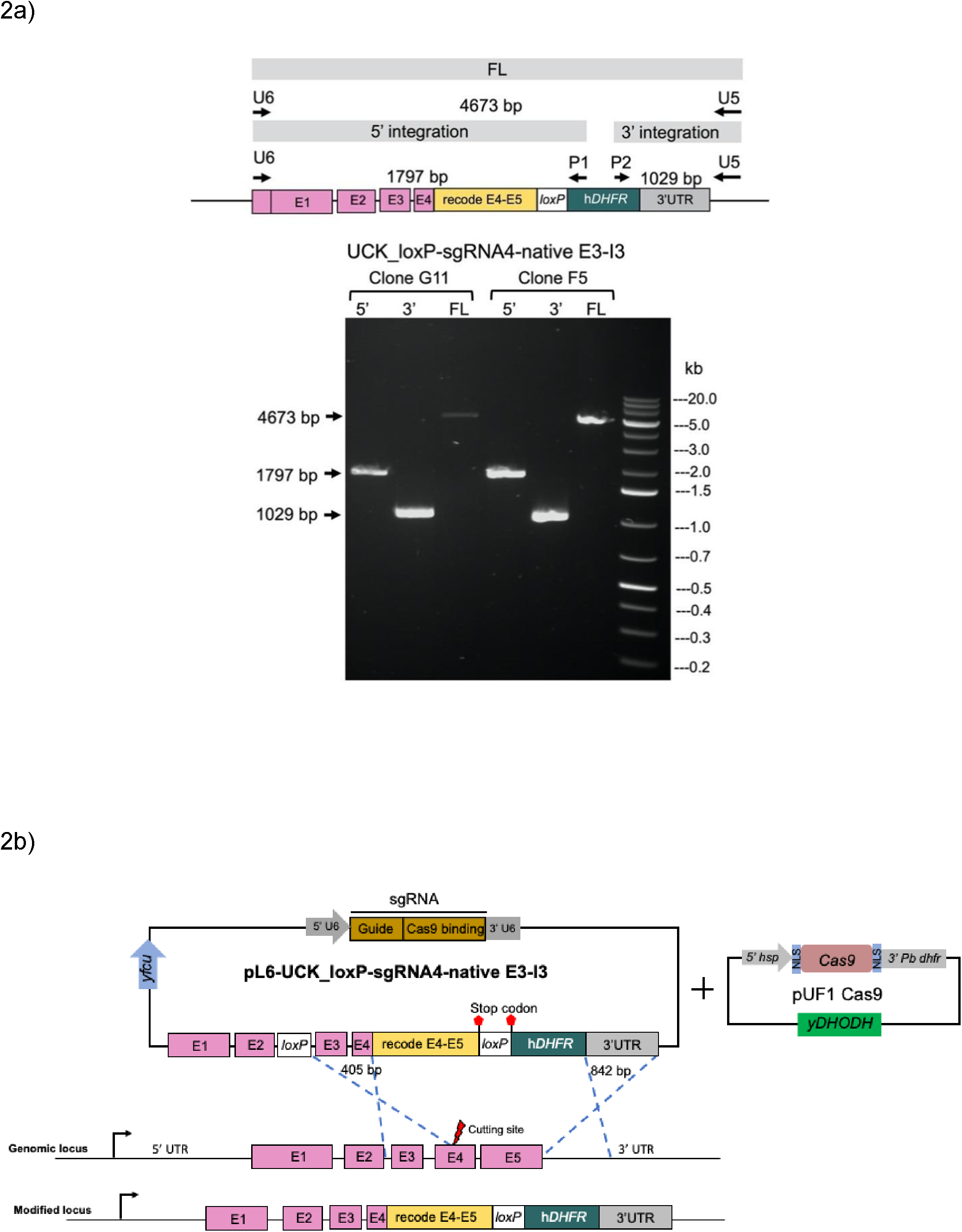
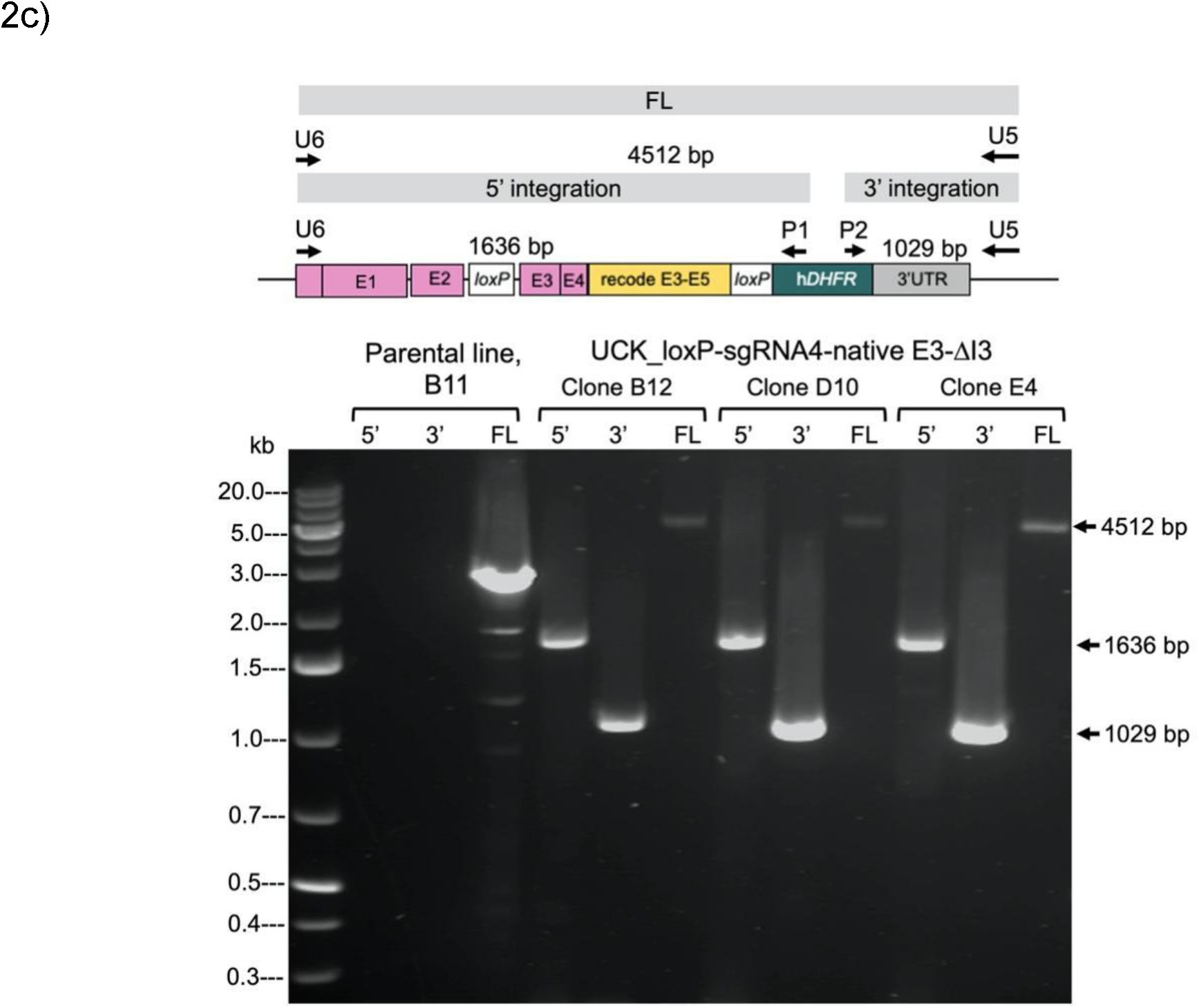
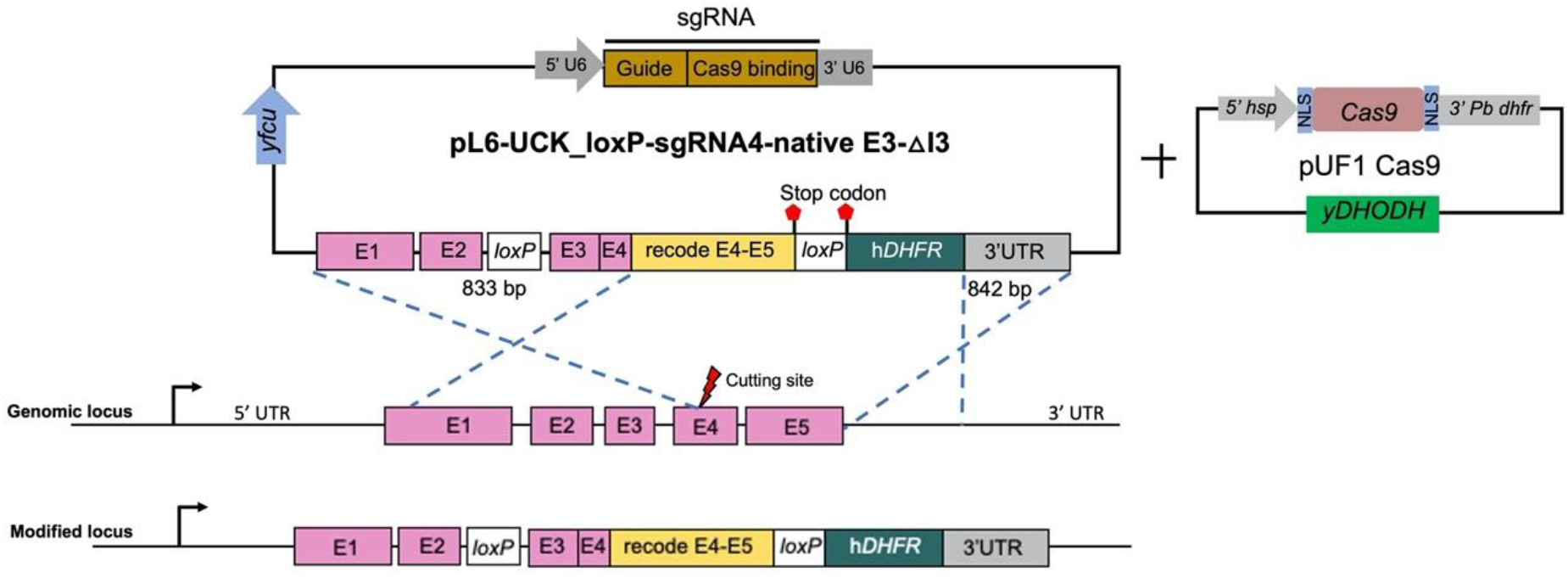
Diagnostic PCR Analysis and Homologous Recombination Scheme. a) Diagnostic PCR analysis of genomic DNA from integrant P. *falciparum* clones (G11 and F5) obtained from transfections using the repair plasmid pL6-UCK_loxP-sgRNA4-native E3-I3. Both clones exhibited positive genotyping for 5’ and 3’ integration as well as the full length (FL) of the modified Pf*UCK* locus. Whereas DNA sequencing analysis of transfectant clone F5 shows the absence of the upstream *loxP* sequence. b) Diagram illustrating a possible homologous recombination event in the transfectant clone F5, resulting in the integration of only the downstream *loxP* site into the target locus. c) Diagnostic PCR analysis of genomic DNA from integrant P. *falciparum* clones (B12, D10, and E4) derived from transfections using the repair plasmid pL6-UCK_loxP-sgRNA4-native E3-ΔI3. These clones exhibited positive genotyping for 5’ and 3’ integration as well as the complete modification of the Pf*UCK* locus. d) Diagram illustrating the predicted homologous recombination event in transfectants of pL6-UCK_loxP-sgRNA4-native E3-ΔI3.

Mutant parasites were generated by transfecting the repair plasmids into the DiCre-expressing P. *falciparum* B11 line (20). Successful modification of the Pf*UCK* gene in the transfected parasite population was confirmed through diagnostic PCR. Subsequent limiting dilution cloning resulted in the generation of parasite clones G11 and F5 using the repair plasmid pL6-UCK_loxP-sgRNA4-native E3-I3 (Figure 2a). Sequencing analysis of pL6-UCK_loxP-sgRNA4-native E3-I3 transfectant clone F5 revealed integration of only the downstream *loxP* site into the target locus, while the upstream *loxP* site was absent, potentially indicating alternative recombination during the repair process. Since this repair plasmid was designed to retain the native exon 3 and intron 3, a sequence of 405 bp between native exons 3 and 4 was preserved, which is sufficient for homologous recombination. Previous studies have shown that homologous regions of >250-1000 bp are adequate for homology-directed repair in *P. falciparum* (46). Therefore, the 405 bp segment, situated close to the Cas nuclease cleavage site, exhibits potential for homologous recombination, as illustrated in Figure 2b.

The transfectant parasites of pL6-UCK_loxP-sgRNA4-native E3-ΔI3, isolated as clones B12, D10, and E4, exhibited positive genotyping for 5’ and 3’ integration and complete modification of the Pf*UCK* locus (Figure 2c). DNA sequencing of transfectant clone E4 confirmed the successful integration of both upstream and downstream *loxP* sequences at the target Pf*UCK* locus. This indicates the generation of transgenic parasites through the expected homologous recombination, as depicted in Figure 2d.

### 3.4. Generation of UCK knockout mutants

The insertion of the two *loxP* sites allows truncation of *UCK* by inducing DiCre expression. Hence, the clones with complete integration (referred to as UCK-clone B12, D10, and E4), were induced to delete the majority of the *UCK* gene (Figure 2c & 2d). As a control, the clone F5 that harbors a single *loxP* site positioned at the end of the *UCK* gene was used. This parasite clone was designated as UCK-control F5 and served as a control for unexcised parasites following rapamycin (RAP) treatment.

To induce *UCK* gene truncation, tightly synchronized ring-stage transgenic parasite clones were subjected to treatment with either 50 nM RAP or a mock treatment using 0.1% DMSO for 24 hours. Expected site-specific recombination between the introduced *loxP* sites in the modified Pf*UCK* locus of clones B12, D10, and E4 occurred upon RAP treatment, resulting in the reconstitution of a single *loxP*-containing parasite (excised locus; Figure 3). Diagnostic PCR results revealed a 1,636 bp band in DMSO-treated samples and an 835 bp band in RAP-treated samples, confirming the anticipated excision event. Conversely, only unexcised bands were amplified as expected in parasite UCK-control F5. Notably, non-excised DNA was detected in UCK clone B12.

**Figure 3.**
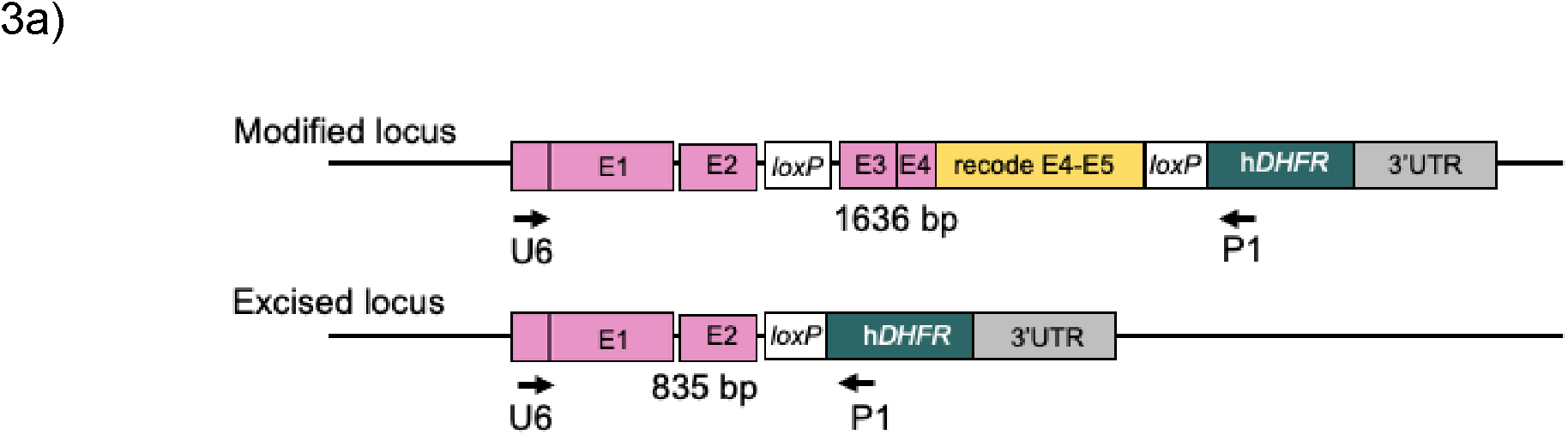
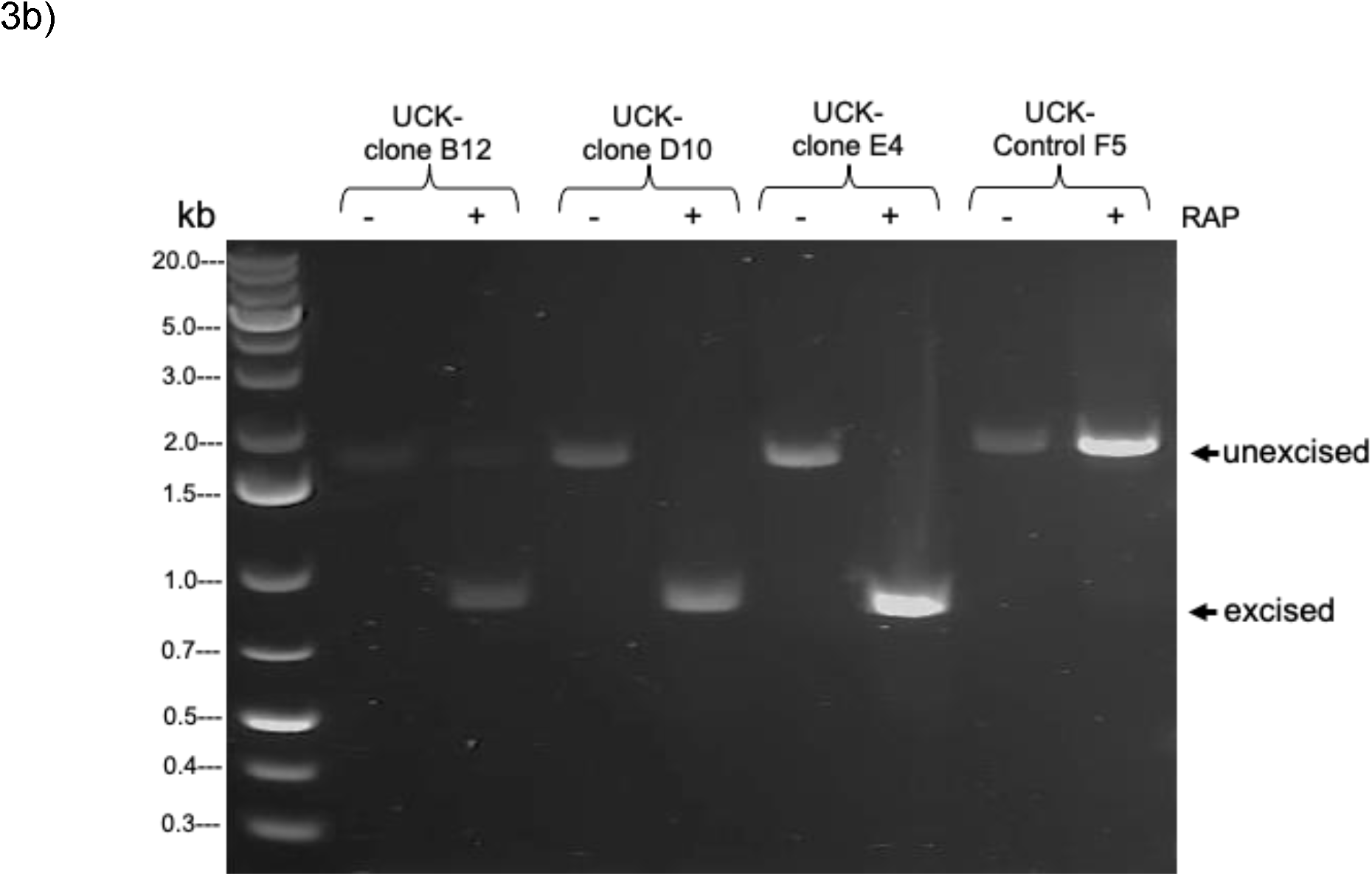
DiCre-mediated conditional disruption of Pf*UCK*. a) An illustration of the primers specifically designed for the detection of Pf*UCK* truncated parasites. b) PCR products of mutants following rapamycin induction of cre/lox deletion. At 42 hours post-RAP treatment, 200 µL of parasite cultures were harvested for gDNA extraction and PCR analysis of DiCre-mediated excision of the *loxP*-floxed DNA. The expected sizes of the PCR amplicons, including unexcised and excised bands, are indicated by arrows.

### 3.5. Phenotype of parasite UCK induced knockouts

Since maximal expression of *UCK* is observed in trophozoites and persists into the schizont stage (18), trophozoite stage parasites were selected as the source for studying gene expression levels, comparing untreated and RAP-treated parasites. Levels of Pf*UCK* mRNA in UCK-clones B12, D10, and E4 were significantly reduced (approximately 45–230 fold) following RAP treatment (incubation with 50 nM RAP for 24 hours), based on qRT-PCR (Figure 4). However, Pf*UCK* mRNA levels in the control F5 were unaffected by RAP treatment.

**Figure 4.**
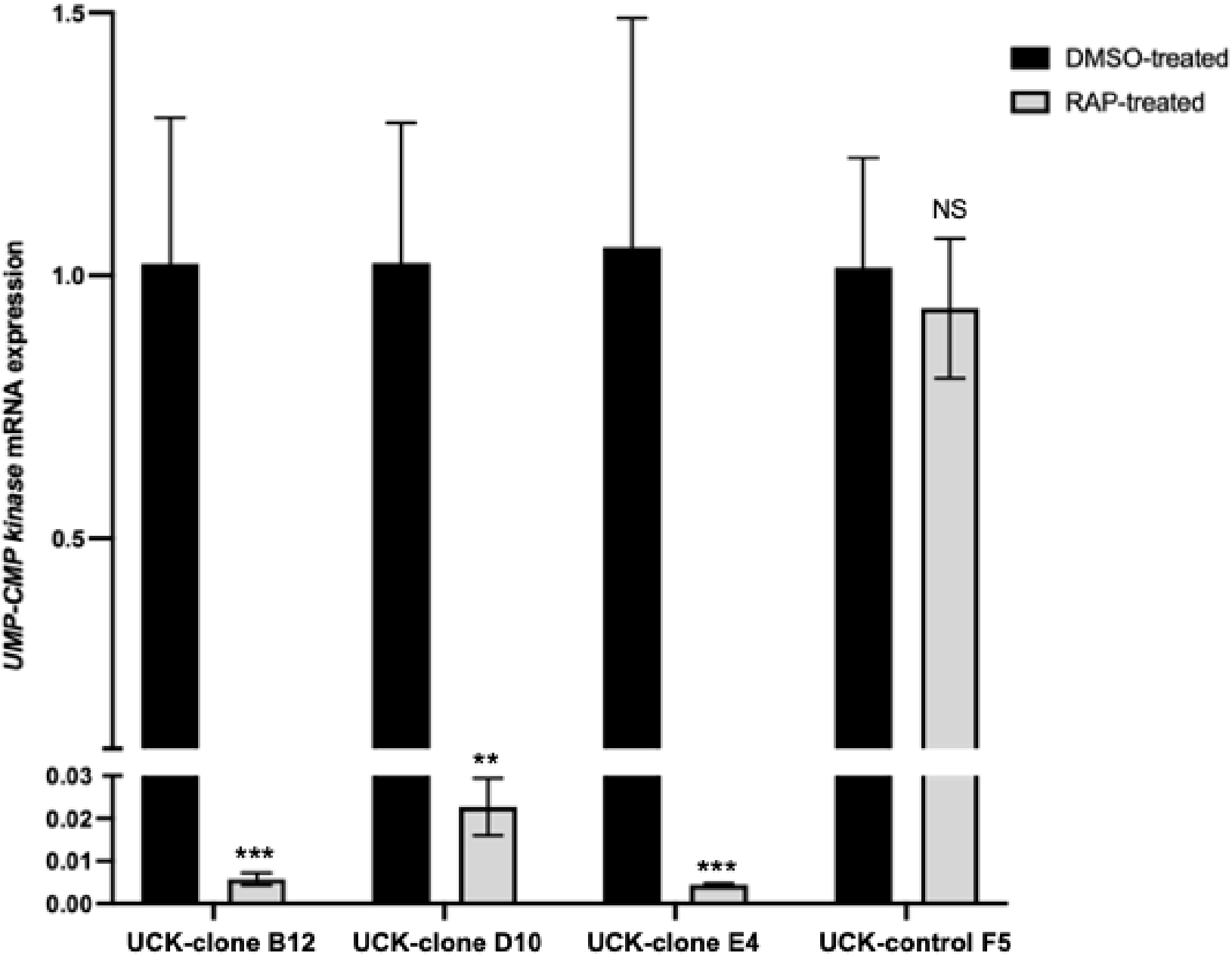
Analysis of Pf*UCK* mRNA expression in response to RAP.

RNA from rapamycin treated parasites or placebo control was isolated from the transgenic clones and subjected to RT-qPCR for Pf*UCK*. Transgenic parasite F5 (bearing a single *loxP* site) was used as the control for Pf*UCK* gene expression. The levels of Pf*UCK* mRNA in RAP-treated cultures relative to DMSO-treated cultures were determined by RT-qPCR using the 2^-ΔΔCT^ method by normalising to *SSU* rRNA. Error bars represent the standard error of the mean across three replicates. Statistical analysis using two-tailed Student’s *t-*tests was performed to assess the significance of changes in mRNA levels as paired samples. NS denotes not significant, and a *P*<0.05 was considered statistically significant with ***, *P*<0.0001; **, *P*<0.001.

### 3.6. Dependence of parasite growth on UCK

To assess the growth effects of disruption of Pf*UCK*, a growth assay based on FACS of SYBR green-labeled transgenic parasites was performed over two cycles of intraerythrocytic development for parasites treated with DMSO (control) compared with RAP. Four days following RAP treatment the parasitaemia levels for *UCK* deleted parasites were significantly less than the controls (Figure 5). Decreased growth was slightly observable after the first cycle of replication following RAP treatment (i.e. clones B12 and E4). In contrast, growth of the parasite clone carrying a single *loxP* site was unaffected by RAP treatment. For parasite generations cycle 0 (day 2) and cycle 1 (day 4), parasitaemia levels of UCK-clones B12, D10, and E4 did not increase, indicating that *UCK* is essential for growth and replication of intraerythrocytic parasites.

**Figure 5.**
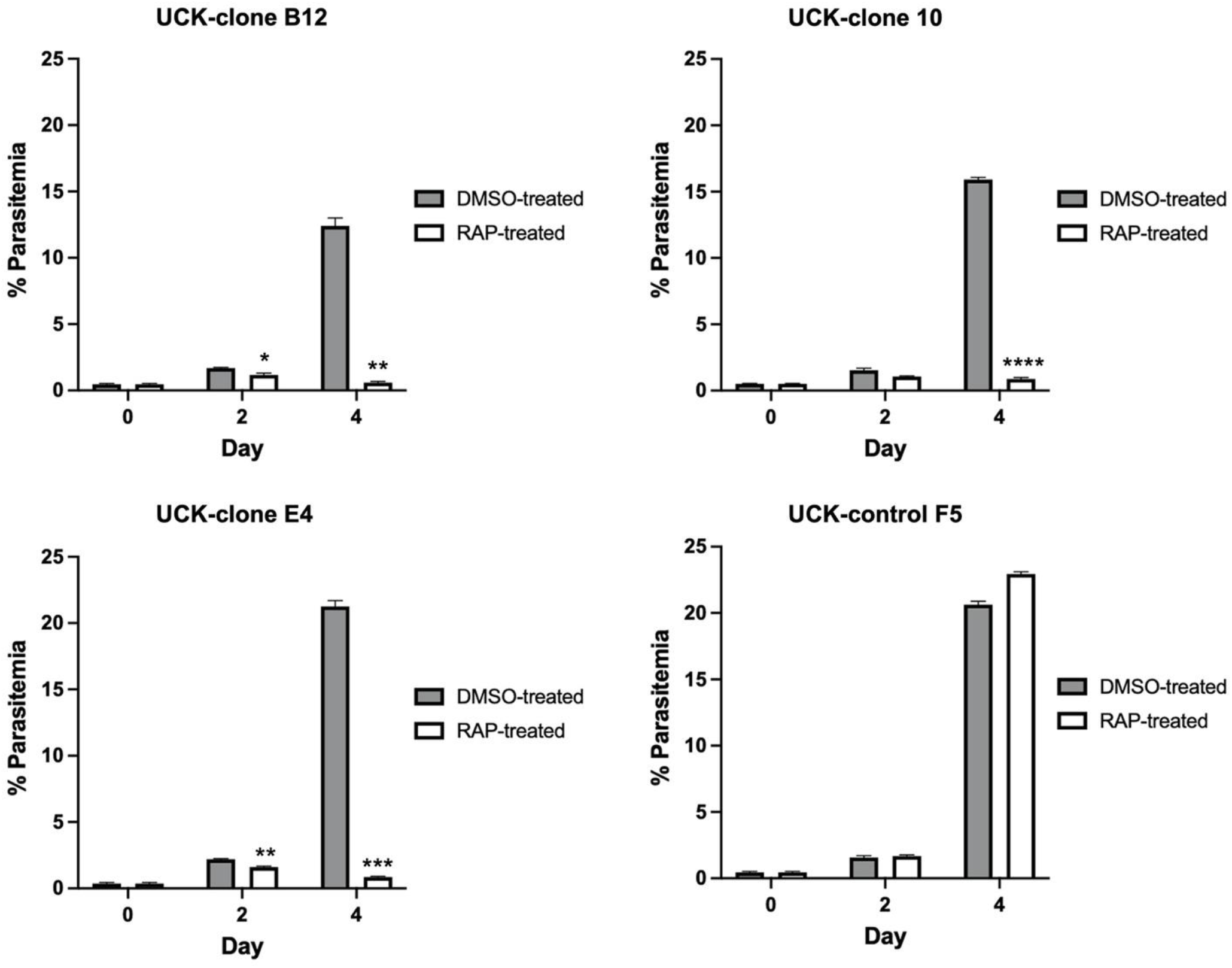
Effect of induced Pf*UCK* disruption on parasite growth. Parasitaemia values were determined by FACS (each data point represents the mean ± standard error across triplicate counts (100,000 events). Statistical analysis of parasitaemia in the RAP-treated parasite culture relative to the control (DMSO-treated) culture was conducted using two-way ANOVA (Bonferroni’s multiple comparison test) with ****, *P*<0.0001; ***, *P*<0.001; **, *P*<0.005; *, *P*<0.05.

### 3.7. Stage-dependent effects of UCK deletion

To assess the effects of *UCK* deletion on parasite development, synchronised ring-stage parasites of transgenic clones UCK-clone B12, UCK-clone D10, and UCK-clone E4 were treated with 50 nM RAP (or placebo 0.1% DMSO) for 24 hours and monitored by microscopy. The UCK-control F5 was used as the control for unexcised parasites. Parasite development was monitored using Giemsa-stained smears in comparison to both DMSO-treated parasites and UCK-control F5 control parasites at 40, 55, and 92 hours post-invasion (Figure 6). The first cycle progressed normally with parasite development from rings to schizonts appearing normal at forty hours post-invasion and transgenic parasites cycling to new ring-stage parasites at fifty-five hours. After the next cycle (cycle 1 at 92 hours post invasion), progression of development had ceased for UCK truncated parasites (UCK-clones B12, D10, and E4 RAP-treated parasites) and they were stalled at the trophozoite stage and appeared unhealthy with shrunken trophozoites. The DMSO-treated control cultures and parasite UCK-control F5 progressed normally with healthy, normal late schizonts and early ring-stage parasites visible. These results support that *UCK* ablation causes a severe defect in parasite development that becomes apparent after the first growth cycle. Hence *UCK* is essential for growth and replication of *P. falciparum*.

**Figure 6.**
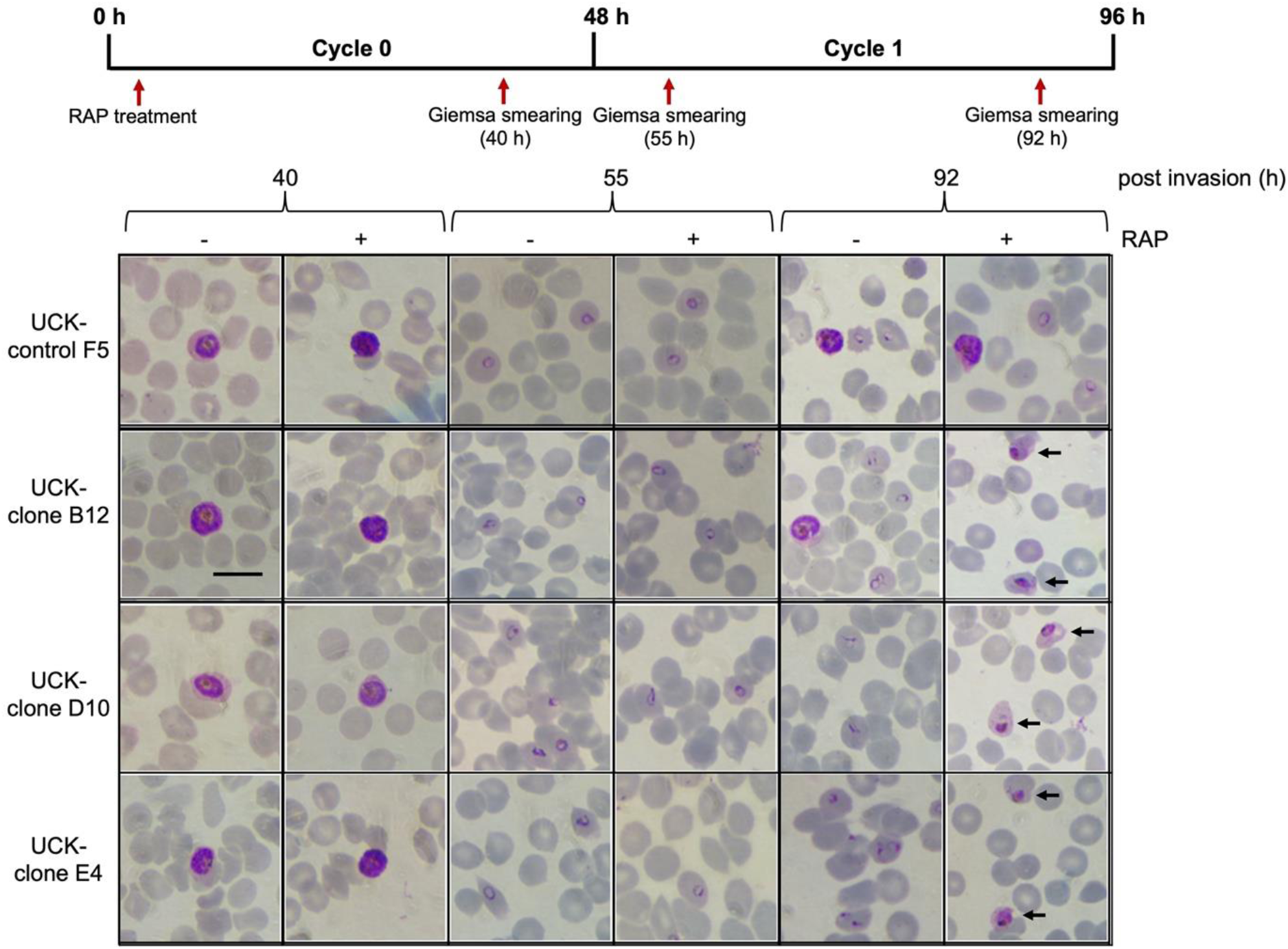
Effect of Pf*UCK* truncation on parasite development based on microscopic examination.

Giemsa-stained thin blood smearing shows the morphology of the truncated compared to the control (DMSO-treated) parasites. Unhealthy trophozoites of disrupted parasite UCK-clones B12, D10, and E4 (RAP-treated) were detected in the following cycle (at 96 hours-post invasion) The time-course of sampling of the cultures is shown on top. Scale bar equals 10 µm.

### 3.8. Biochemical properties of PfUCK

Recombinant PfUCK was expressed and purified for biochemical analyses. To prevent interference from predicted transmembrane domains, the construct had the amino-terminal 23 amino acids truncated. Therefore, the recombinant enzyme PfUCK-Δ23 was utilized in all biochemical analyses.

The kinetic parameters of PfUCK were compared with its human ortholog, hUCK, and the summarized results are presented (Table 3 and Figure 7). Enzyme activity assays, utilizing ATP as a phosphate donor, demonstrated PfUCK’s preference for ribonucleoside monophosphates CMP and UMP as substrates over dCMP. PfUCK exhibited the lowest *K*_m_ for CMP (28 µM), while UMP displayed a 3.9-fold higher *K*_m_ (110 µM). In contrast, the *K*_m_ for dCMP was notably higher at 428 µM compared to CMP and UMP. Similar to that of PfUCK, hUCK displayed a *K*_m_ for CMP (24.8 µM), while its *K*_m_ for UMP was 2.3-fold greater than PfUCK.

**Table 3.** PfUCK kinetic parameters.

| Phosphate<br>acceptor | PfUCK | hUCK | hUCK* |
| --- | --- | --- | --- |
| | $K_m$ ($\mu\text{M}$) | $K_m$ ($\mu\text{M}$) | $K_m$ ($\mu\text{M}$) |
| UMP | $110 \pm 14$ | $47.7 \pm 3.2$ | $45 \pm 10$ |
| CMP | $28 \pm 9$ | $24.8 \pm 2.6$ | $20 \pm 5$ |
| dCMP | $428 \pm 11$ | nd | $900 \pm 100$ |

hUCK\* was obtained from (47) based on the activity assay was carried out using spectrophotometer in the coupled reaction with pyruvate kinase/lactate dehydrogenase and used 10 mM DTT in the assay reaction.

**Figure 7.**
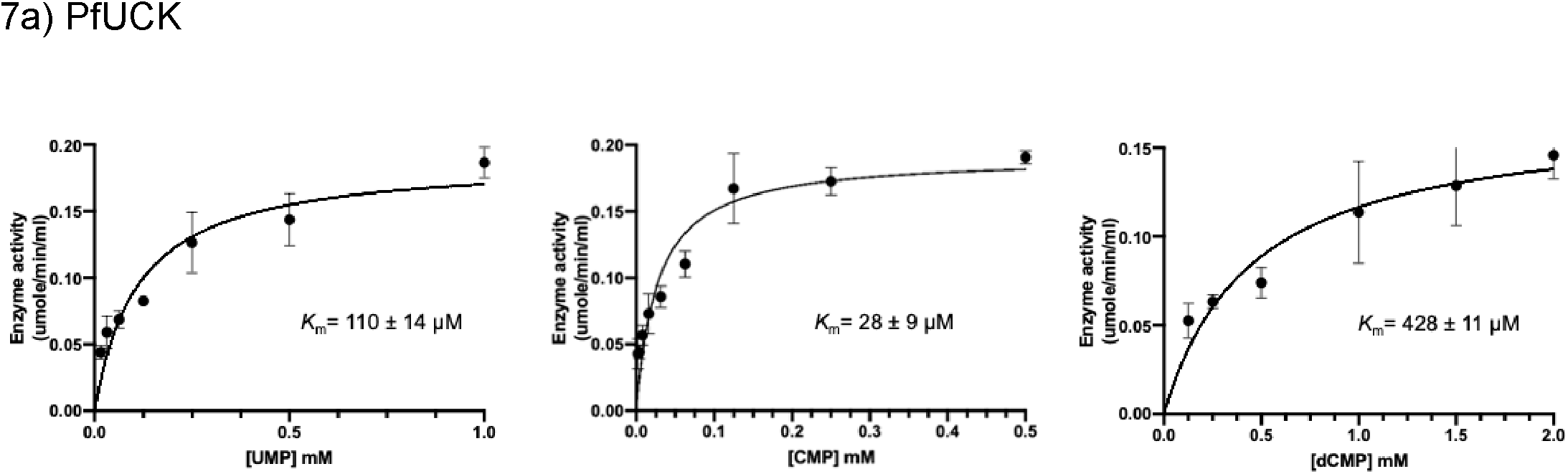
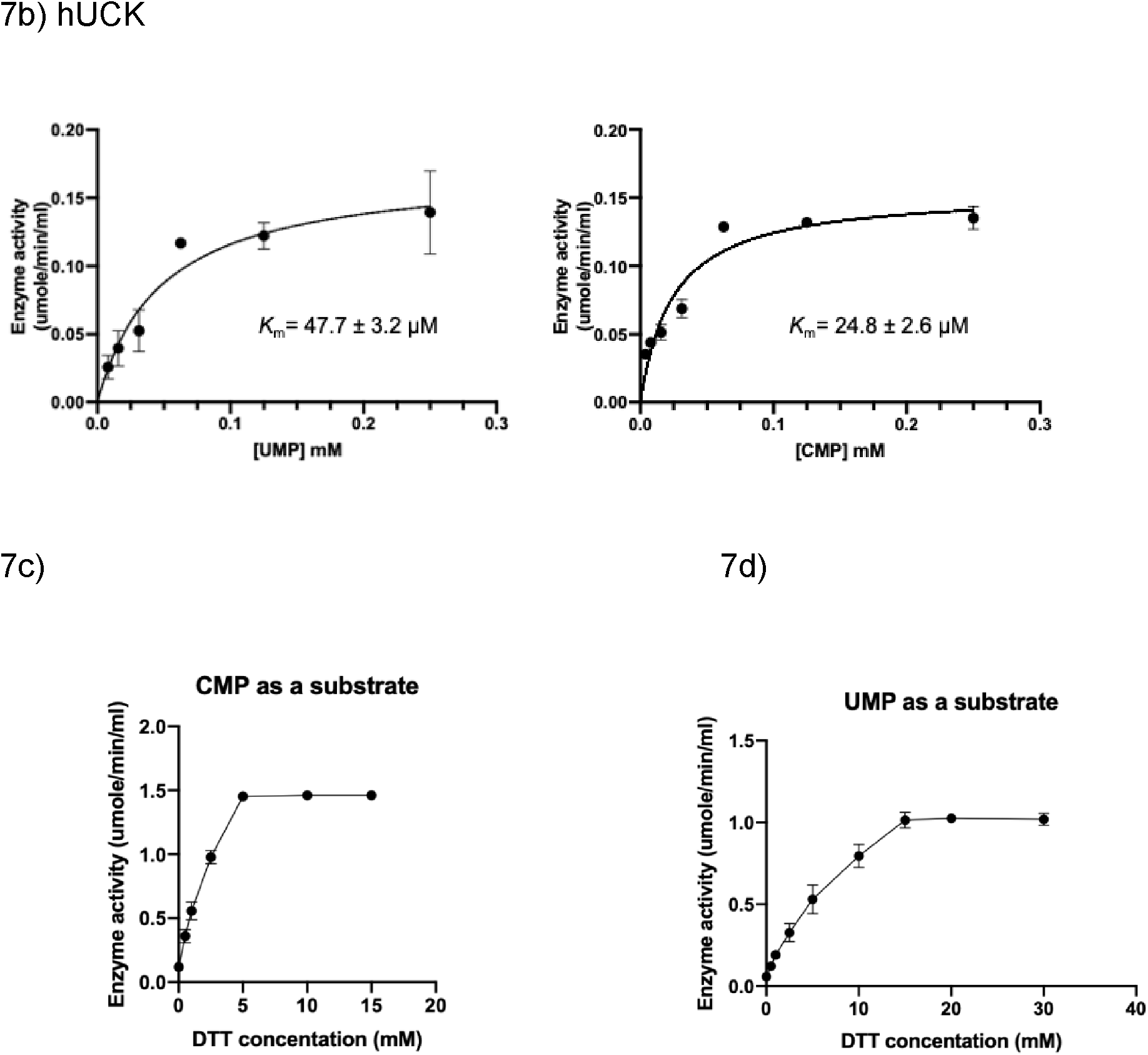
Substrate affinity and reducing dependence of PfUCK.

The substrate binding affinity of hUCK observed in this study are similar to those previously reported by Pasti (47). Regarding substrate preferences, this investigation revealed that ribonucleoside monophosphates (CMP and UMP) are favoured substrates for both human and *Plasmodium* parasites, compared to deoxyribonucleoside monophosphates (dCMP). The most preferred substrate for both hUCK and PfUCK was CMP.

We investigated the effect of reducing agents on PfUCK activity, illustrated in Figure 7c and 7d, as it was previously reported that reducing agents activate UCK kinase activity (48). As found with hUCK, PfUCK activity was enhanced in response to reducing conditions.

Enzyme activities were assessed by varying substrate concentrations, and *K*_m_ values were determined using the Michaelis-Menten equation in GraphPad Prism software (v = Vmax[S] / *K*_m_ +[S]), where v represents the reaction rate, [S] denotes the substrate concentration, and *K*_m_ signifies the Michaelis constant. Graphs depicting *K*_m_ determination for PfUCK (a) and hUCK (b) are shown. Additionally, purified protein under a nonreducing condition was tested its activity by varying DTT concentrations, using CMP (c) and UMP (d) as substrates.

### 3.9. Inhibitor development for PfUCK

A set of compounds was selected to inhibit UCK activity based on in silico screening. Currently, there are no known UCK inhibitors. A homology model of PfUCK was developed based on the structure of UMP/CMP kinase from Dictyostelium discoideum (2UKD). The strategy for the inhibitor design was to identify covalent inhibitors of the enzyme (targeted towards Cys139). Virtual screening was performed using the CovDock tool within Glide (49) focused on modulation of Cys139. Covalent virtual libraries were selected from a number of vendors to include a range of electrophilic warheads thought suitable for cysteine modification, including acrylamide, chloroacetamide and benzoyl acetonitrile. Screening results for acrylamide and chloroacetamides against PfUCK and human UCK (hUCK) are summarized in Table 2. Among these compounds, the pyrimidin-4-one scaffold was present within three hits accommodated both within chloroacetamide and acrylamide warheads and exhibited significant inhibition activity against PfUCK, with a *K*_i_ value in the single micromolar range. Unfortunately, C43 and C21 also inhibited hUCK activity, with a similar *K*_i_ value in the same range. Hence, further refinement and selectivity optimization were needed for UCK inhibitors to avoid possible cytotoxicity.

Amongst the benzoylacetonitrile derivatives (Table 5), several exhibited micromolar level inhibition of PfUCK highlighting C38, which contains a fluoro-group at the meta-position which increases the reactivity resulting the most potent inhibitor with a *K*_i_ value of 1.66 ± 0.27 µM. Based on the structure-activity relationship, the lack of inhibition by C69 is inexplicable. C38, C39 (ortho-chloro derivative), and C7 (meta-chloro derivative) exhibited selectivity for PfUCK with little or no inhibition of hUCK. These results highlight the crucial role of specific substituents in determining both inhibitory potency and selectivity against PfUCK.

**Table 4.**
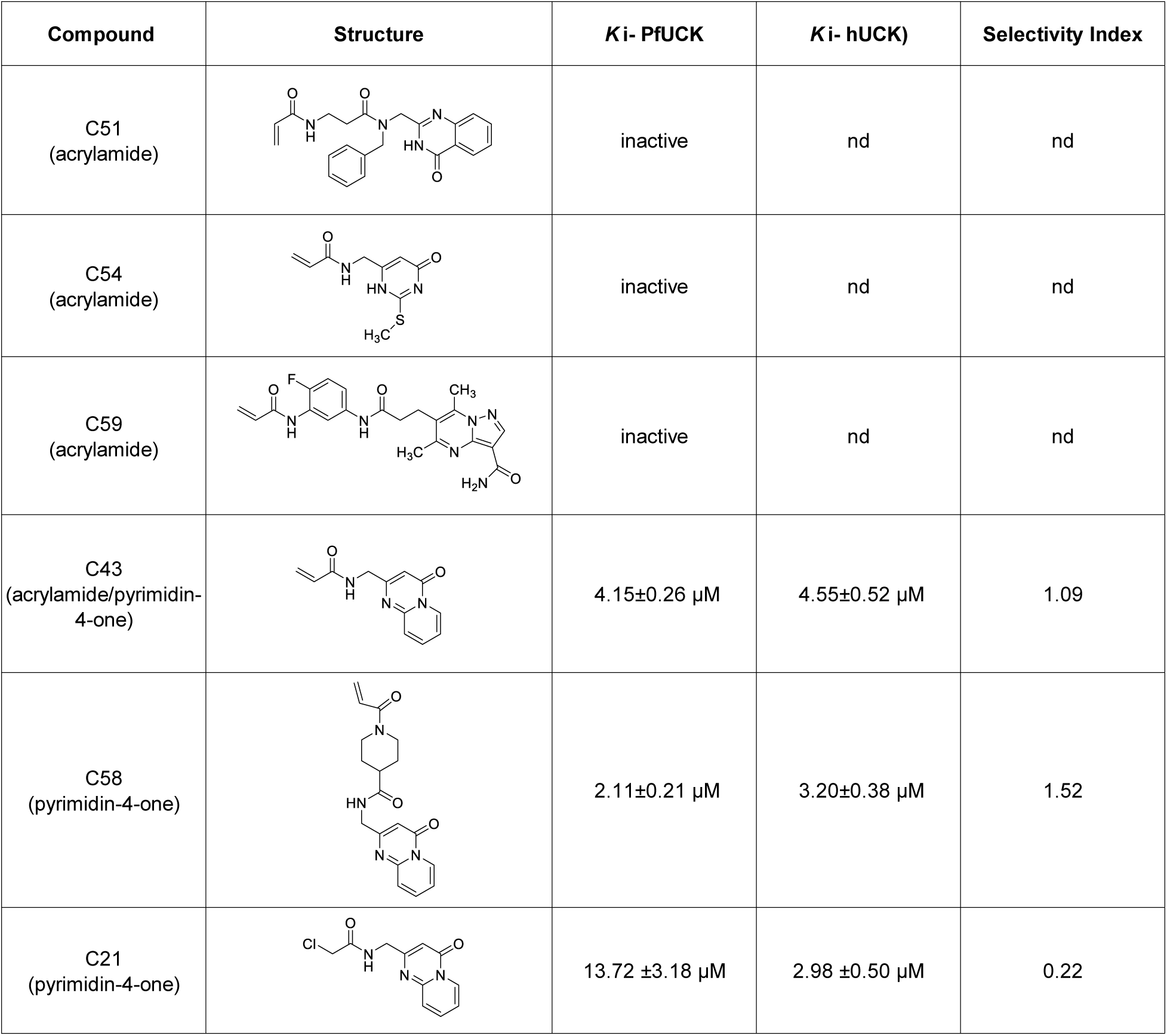
Screening acrylamide and chloroacetamide compounds against PfUCK and hUCK.

| Compound | Structure | K<sub>i</sub>- PfUCK | K<sub>i</sub>- hUCK) | Selectivity Index |
| --- | --- | --- | --- | --- |
| C51<br>(acrylamide) |  | inactive | nd | nd |
| C54<br>(acrylamide) |  | inactive | nd | nd |
| C59<br>(acrylamide) |  | inactive | nd | nd |
| C43<br>(acrylamide/pyrimidin-4-one) |  | 4.15±0.26 μM | 4.55±0.52 μM | 1.09 |
| C58<br>(pyrimidin-4-one) |  | 2.11±0.21 μM | 3.20±0.38 μM | 1.52 |
| C21<br>(pyrimidin-4-one) |  | 13.72 ±3.18 μM | 2.98 ±0.50 μM | 0.22 |

**Table 5.**
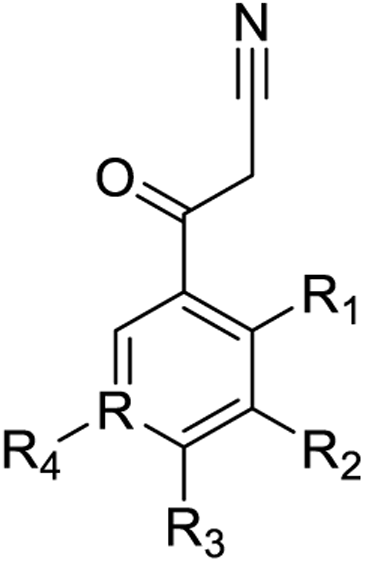
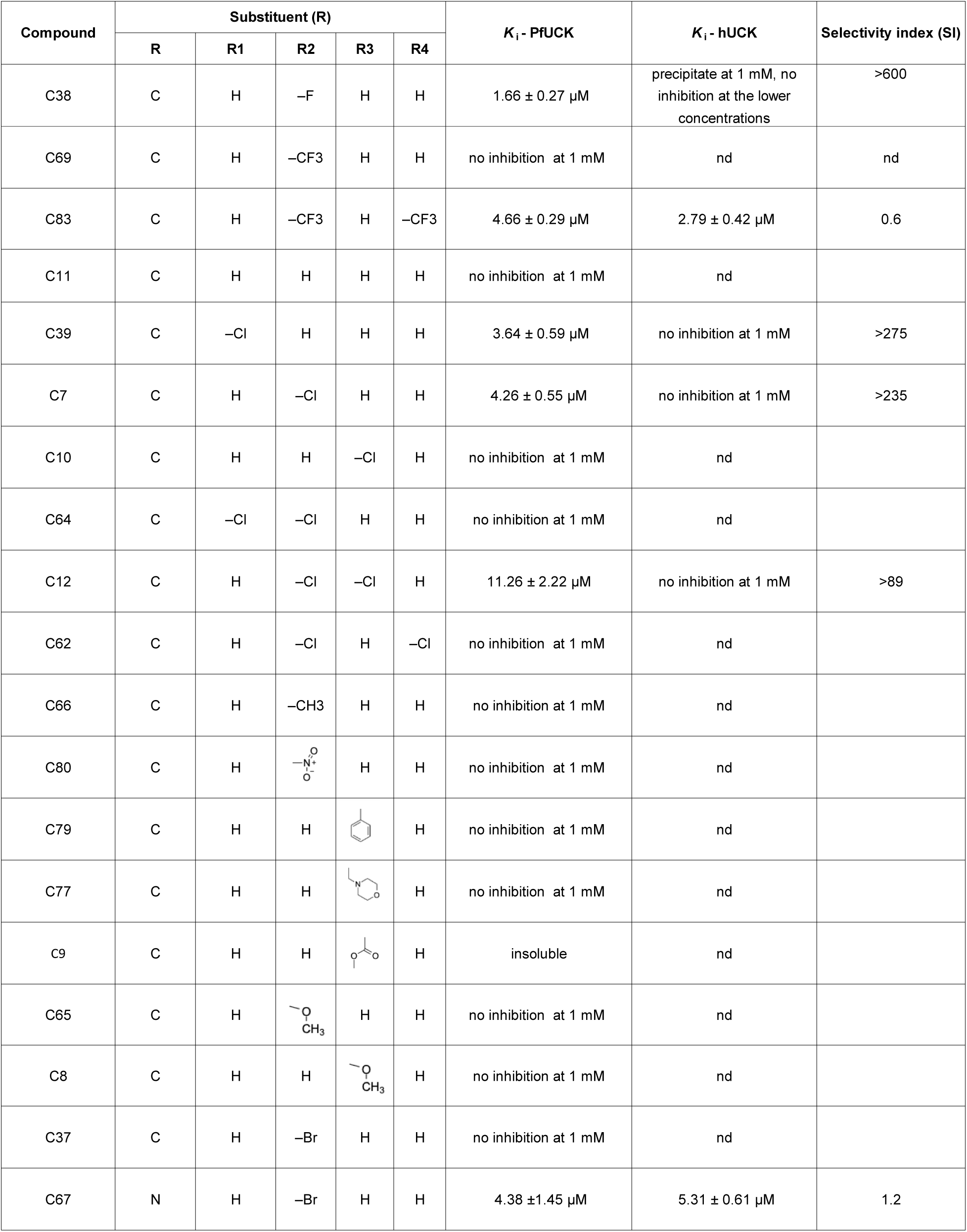
Screening benzoylacetonitrile derivative compounds against PfUCK and hUCK.

Among the compounds tested, those that inhibited PfUCK were evaluated for their effect on *P. falciparum* growth (IC_50_). The only inhibitor with significant antiparasitic activity was the pyrimidin-4-one derivative C21 with an IC_50_ value of 15.64 ± 1.13 µM (Table 4 includes acrylamide and pyrimidin-4-one tested compounds). Regarding the benzoylacetonitrile derivatives, four of them displayed weak inhibitory activities against *P. falciparum* with IC_50_ values exceeding 310 µM (Supplemental Table 4).

**Table 6.** Antiparasitic activities of PfUCK inhibitors.

| Compound | Structure | IC<sub>50</sub> - *P. falciparum* (3D7) |
| --- | --- | --- |
| C43 |  | > 1 mM |
| C58 |  | > 1 mM |
| C21 |  | 15.64 ± 1.13 µM |

## 4. Discussion

This project aimed to test the essentiality and druggability of PfUCK that was predicted from *in silico* metabolic modelling of *P. falciparum* (50). The model identified a set of proteins that were predicted to be essential and hence potential drug targets. Utilising a CRISPR/Cas9-based approach with inducible gene deletion, UCK in *P. falciparum* was found to be essential for growth and progression through the developmental stages. Intriguingly, the phenotypic effect was not immediate. Dysmorphic trophozoites with poor staining were not observed in RAP-treated cultures until the end of cycle 1 (Day 4) coinciding with decreased growth. These results support that *UCK* ablation severely impacts parasite development, arresting them in the trophozoite stage of the erythrocytic cycle. The time delay in phenotype following conditional knockout of *UCK* could be explained by a delay between RAP exposure and start of excision of the targeted DNA sequence or a factor of this enzyme. According to the generation of conditional gene knockouts in *P. falciparum* using the DiCre recombinase system published by (51), RAP concentrations, exposure times, and timing of excisions after adding RAP during early ring parasites can impact excision. They found that excision could be detected significantly after 20 hours of RAP treatment, and it took more than 30 hours to achieve the maximum excision levels. Alternatively, remaining levels of the enzyme were sufficient until rounds of replication diluted the enzyme to threshold levels. For PfUCK, mRNA is expressed throughout the erythrocytic lifecycle. However, the highest expression is found in trophozoites and continues into schizont stage during exponential replication within the erythrocytes. High expression of *UCK* is also detected in the early ring stage before declining in the mid-ring and late-ring parasites (52). Overall, the mRNA expression profile of *UCK* implies that UCK production is maintained throughout the erythrocytic stage. Thus, the phenotype of UCK using the conditional DiCre-mediated excision could require one or more cycles to be observed.

Both human and parasite UCK orthologues exhibited similar substrate preferences, with CMP and UMP being favored substrates, while dCMP was poorly phosphorylated. Further, a reducing agent enhanced PfUCK activity similar to that observed for hUCK and pig UCK (here and (53) and (48).The sensitivity to the reducing agent fits well with the conservation of cysteine residues in PfUCK (Cys298 and Cys307) with human and pig UCK, that are also responsive to reduction. These contrast *Entamoeba histolytica* UCK that lacks these conserved cysteine residues and is not responsive to reducing agents (54). It has been proposed that the presence of cysteine residues in kinases and phosphatases can influence redox regulation within cells although the concentrations used here are not physiological (55). Notably, in humans, physiological concentrations of thioredoxin have been demonstrated to fully activate kinase activity (48). Consequently, a deeper investigation into the influence of physiological thioredoxin concentrations are warranted. This property may highlight how valuable UCK may prove as a target because of the high sensitivity of the parasite to oxidant levels and redox balance. Indeed, this finding may be advantageous in possible combination therapies with artemisinin or chloroquine that disrupt the parasite’s redox system.

This study identified, for the first time, micromolar level range inhibitors of PfUCK. Among those, reactive site inhibitors that were tested found that the acrylamide-warhead compound C43 exhibited PfUCK inhibitory activity (*K*_i_ value approximately 4 µM). Our modelling predicted interaction of the inhibitors with Cys139. Confirmation of covalent attachment through kinetic assays or mass spectrometry of the protein-ligand complex is needed (56). Future studies require design of new acrylamide warhead compounds with examination of the conjugation. The benzoylacetonitrile derivative compounds exhibited a diverse spectrum of potencies as inhibitors of PfUCK, along with some compounds displaying limited antiparasitic activity. Future design of PfUCK inhibitors would benefit from crystallographic structure data; with ligands such as compound C38 or C7 to understand protein-ligand binding specificity and aid optimal structure-based design.

Combination therapy is essential for antimalarial chemotherapy due to the rapid development of resistance. Inhibitors of PfUCK are well placed to be combined with antimalarial drugs against pyrimidine biosynthesis, hence, targeting two essential steps because *Plasmodium* relies on *de novo* synthesis for supply of pyrimidines. There is potential for combination with dihydrofolate reductase inhibitors (e.g. pyrimethamine, proguanil) or those developed against the recent target dihydroorotate dehydrogenase. Overall, based on our findings, the assertion of a genome-scale metabolic model based on experimental biomass and experimental flux measurements was founded in an inducible gene knockout demonstrating that UCK is essential in Plasmodium. Further development of inhibitors of this essential enzyme with its redox sensitivity and potential for irreversible inhibition may provide an avenue to new antimalarial development.

---

**Supplementary Figure 1.**
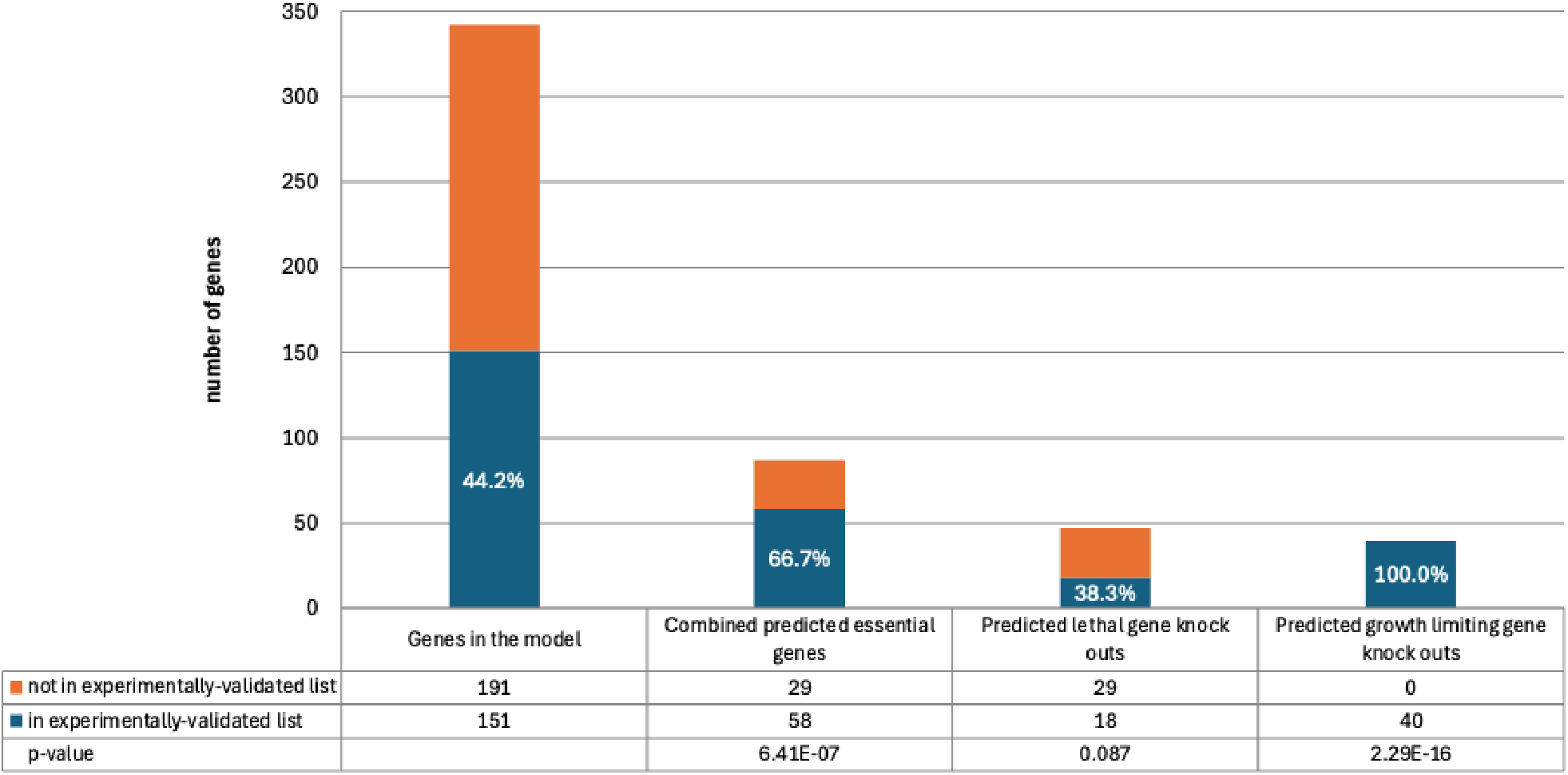
Summary of predicted essential genes. Single gene knockouts with COBRApy were simulated using the schizont stage metabolic model to predict essential genes. The predicted essential genes and their associated reactions were compared with experimentally validated essential reactions. The proportions of true positive predictions (overall, lethal and growth limiting knockouts) were compared against the proportion of genes in the model that are in the gold standard list and the corresponding enrichment p-values were calculated.

**Supplementary Figure 2.**
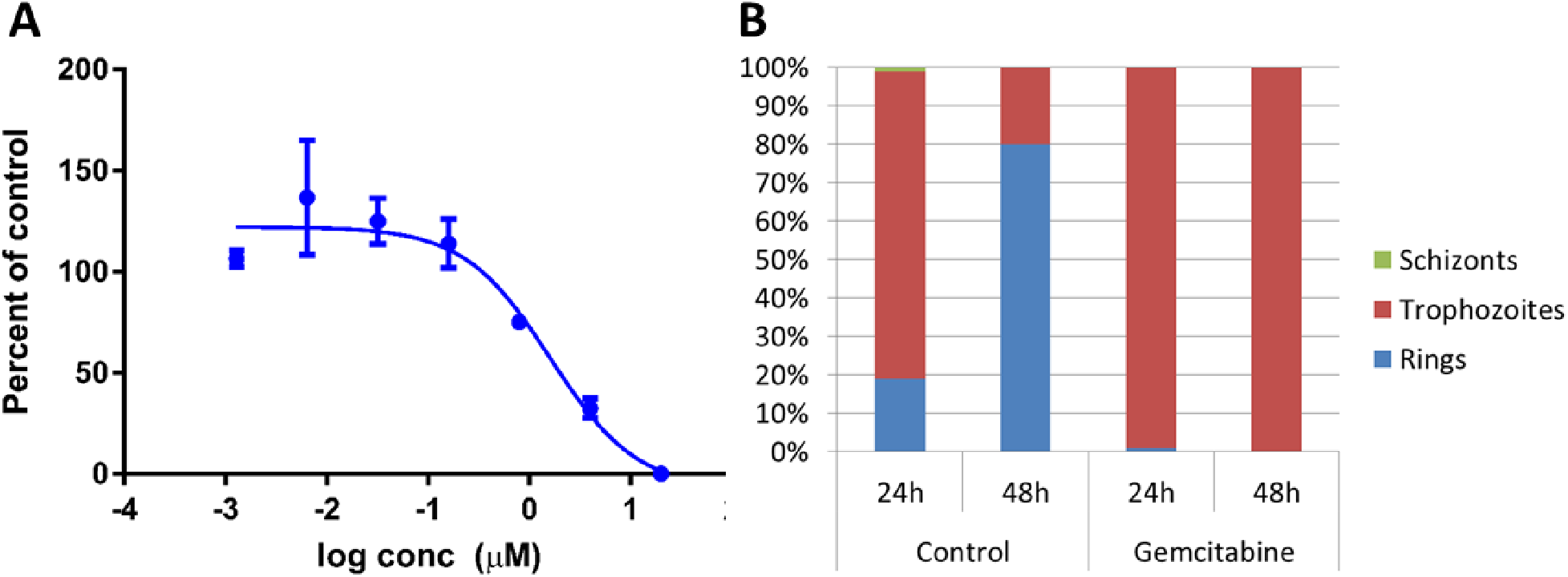
Gemcitabine inhibition of *Plasmodium falciparum* 3D7 growth. (A) Graph of the dose-response to gemcitabine treatment. *P. falciparum* 3D7 cultures were treated with gemcitabine for 48 hours and percent growth relative to untreated controls calculated (n = 3 biological replicates). (B) Graph of developmental stages of parasites following incubation with gemcitabine. Percent of parasites at three stages were calculated in synchronised parasites that were incubated for 24 and 48 hours. In the presence of gemcitabine, late trophozoite stage parasites did not develop into schizonts.

**Supplementary Table 1.**
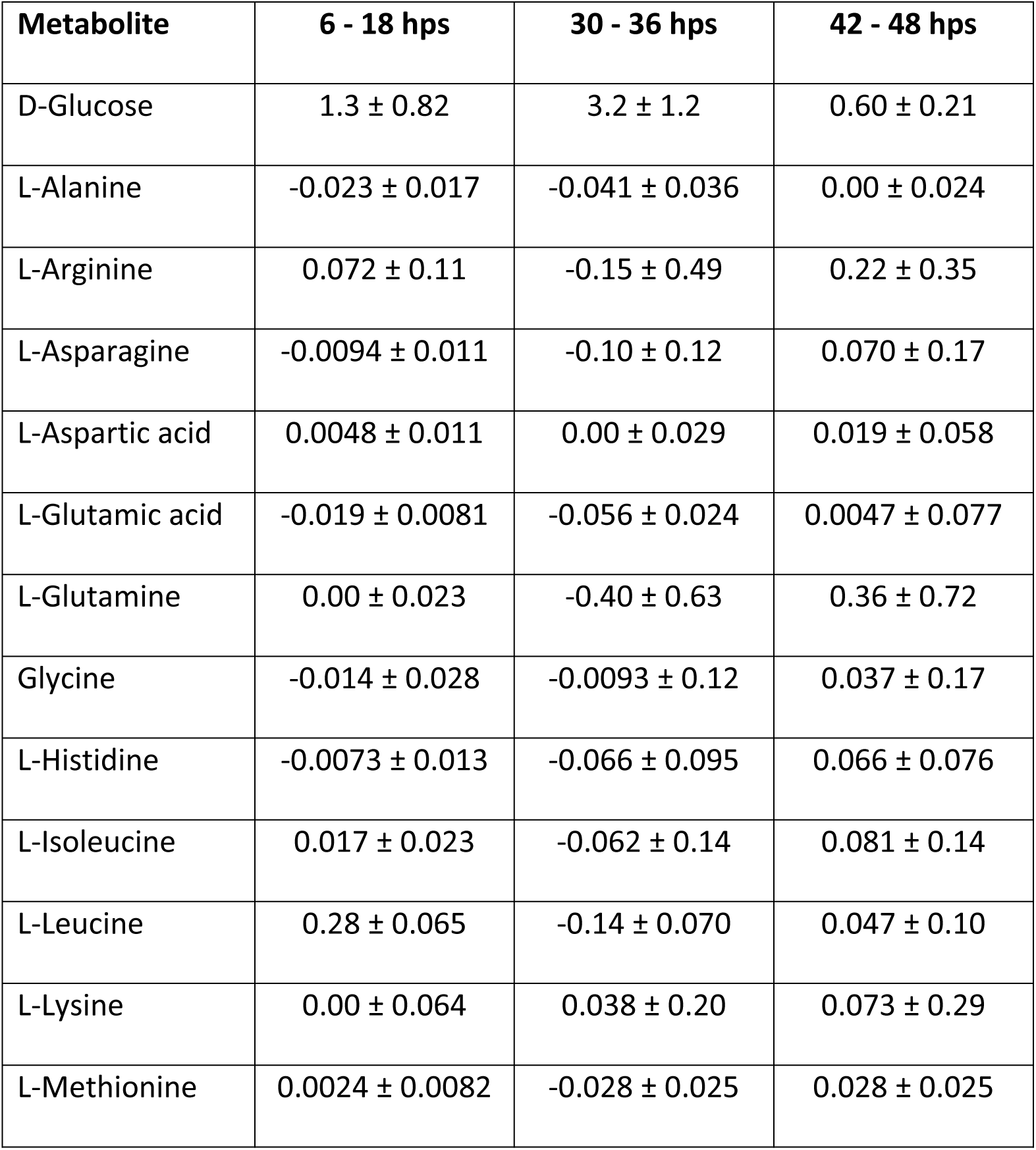
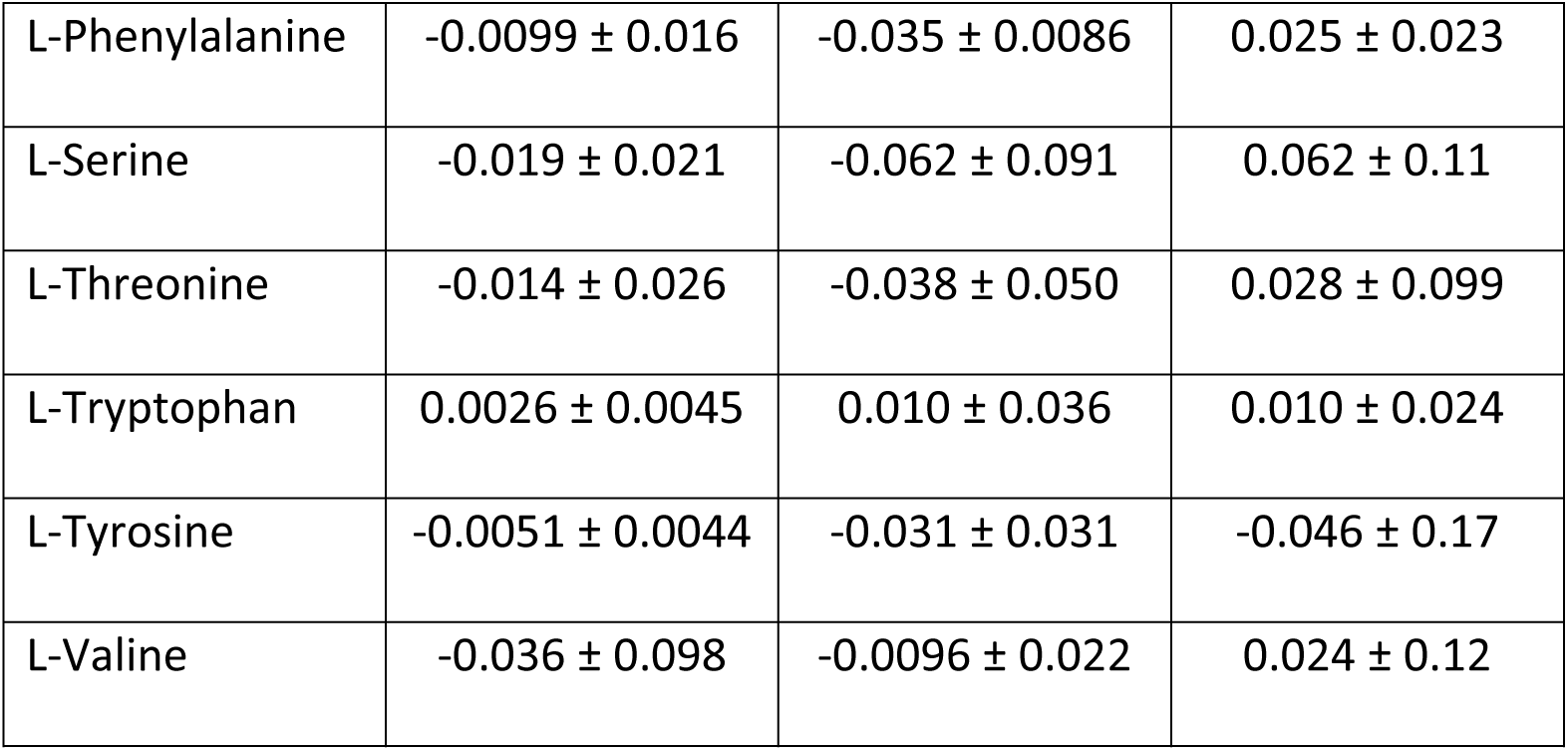
Experimental glucose and amino acid flux measurements. Spent media from synchronised *Plasmodium falciparum* 3D7 cultures were collected at 6, 18, 30, 36, 42 and 48 hours post synchronisation (hps), and glucose and amino acid flux (in mmol/gDW/hr ± SD) were calculated points to represent the early to mid-ring stage, late trophozoite stage and late schizont stage, respectively. A positive flux indicates movement of the metabolite into the infected red blood cell while a negative flux indicates efflux of the metabolite into the media.

**Supplementary Table 2.** List of experimentally validated essential *P. falciparum* genes from published literature, orthologues of essential *P. berghei* genes from PlasmoGEM (16), and combined list of unique genes from both sets. Please see attached Excel file “Supplementary Table 2.xlsx”

**Supplementary Table 3.** HPLC Program.

| **Time (min)** | **Percent Eluent A** | **Percent Eluent B** | **Flow rate (ml/min)** |
| --- | --- | --- | --- |
| 0.0 | 98% | 2% | 0.200 |
| 60.0 | 55% | 45% | 0.200 |
| 60.5 | 98% | 2% | 0.200 |
| 70.5 | 98% | 2% | 0.200 |

**Supplementary Table 4.**
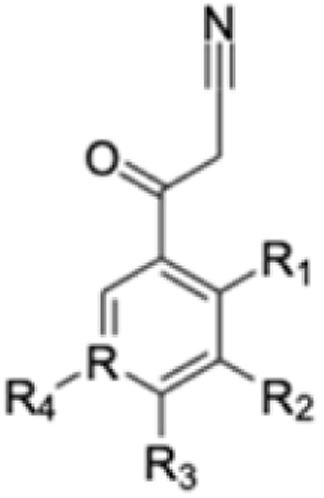
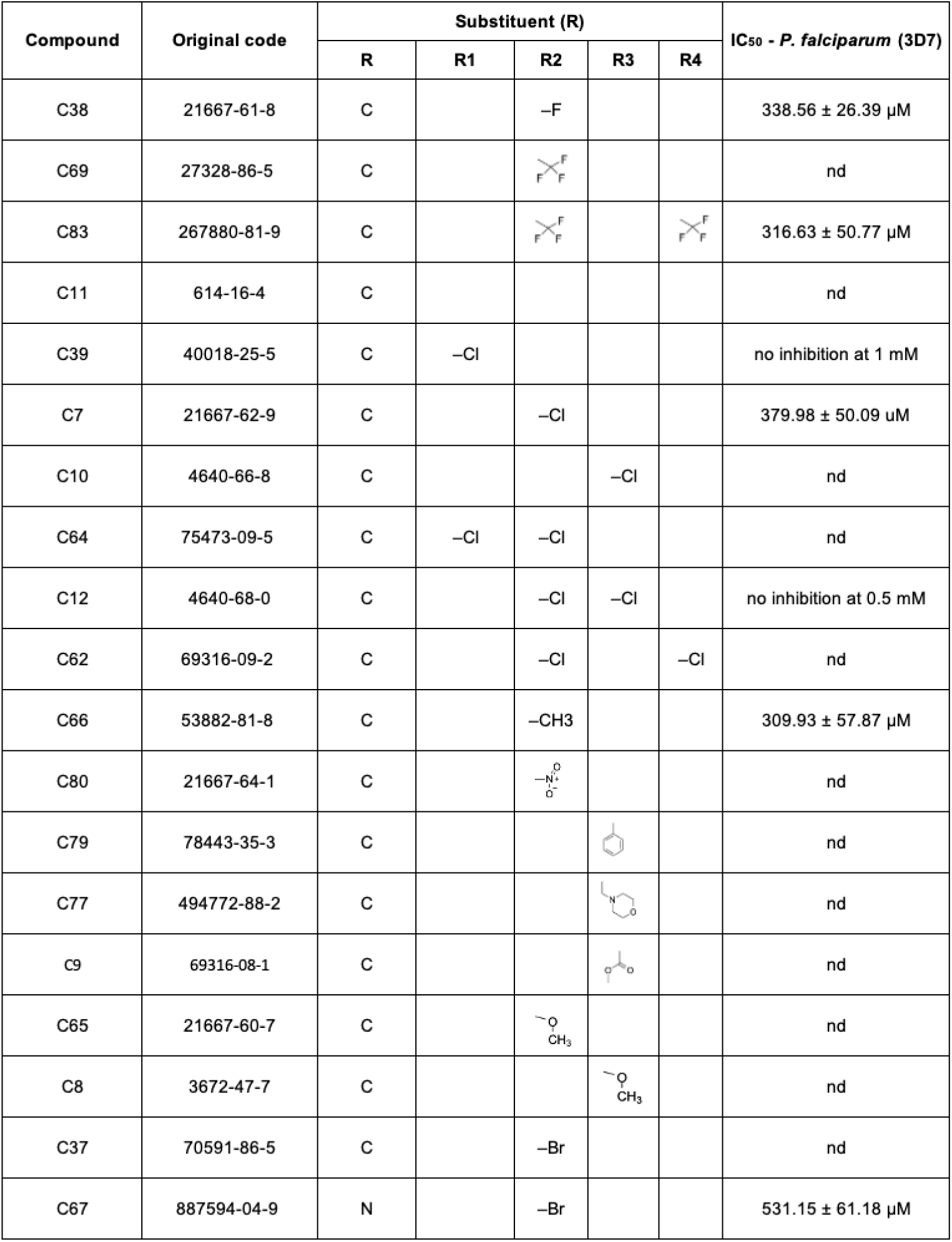
Antiparasitic activities of benzoylacetonitrile derivative compounds.

